# LOX inhibition disrupts a collagen-integrin–MYC axis as a translatable targeting strategy in invasive lobular carcinoma

**DOI:** 10.1101/2025.08.27.672618

**Authors:** Renée L. Flaherty, Flavia Hughes, George Sflomos, Carlos Ronchi, Harriet Kemp, Theo Roumeliotis, Anya A. Nicholas, Giovanna Ambrosini, Amelie Ziehme, Sarah Becker, Yueyun Zhang, Hazel M. Quinn, Laura Battista, Harveena Padda, Solene Pezot, Samuel Jouny, Yanbo Liu, Rachel Brough, Rebecca Marlow, Marjan Iravani, Alicia Okines, Nicholas C. Turner, Athina Stavrodimou, Khalil Zaman, Maryse Fiche, Beatrice A. Howard, Jyoti S. Choudhary, Victoria Sanz-Moreno, Clare M. Isacke, Lara Perryman, Wolfgang Jarolimek, Syed Haider, Christopher J. Lord, Cathrin Brisken

## Abstract

Invasive lobular carcinoma (ILC) accounts for 15% of breast cancers yet lacks specific therapy because ILCs are underrepresented in clinical trials and preclinical models are lacking. We established intraductal xenograft models to test whether the clinical pan-lysyl-oxidase PXS-5505, now in phase trials for myelofibrosis can exploit the collagen-rich matrix dependency created by CDH1 loss. PXS-5505 remodels fibrillar collagen, and halts tumor expansion and metastatic seeding across ER+ and triple negative models without systemic toxicity. Genome-wide CRISPR screens reveal *ITGAV* and *ITGB5* as synthetic lethal partners of *CDH1* and LOX inhibition downregulates their expression together with MYC, NF-κB, and AP-1 transcriptional programmes. Collagen fibre density/alignment, and MYC/AP-1 gene signatures serve as pharmacodynamic readouts of drug activity. These data uncover a tractable ECM-integrin-MYC axis in ILC and nominate PXS-5505, alone or with endocrine therapy, for window of opportunity trials in this understudied breast cancer subtype.

**One Sentence Summary:** Targeting matrix remodelling in ILC inhibits ILC progression and alters multiple molecular endpoints, providing a translatable therapeutic strategy for this understudied subtype that requires better treatments.

## Introduction

Breast cancer is a heterogeneous in both histology and molecular makeup. Approximately 10-15% of cases are invasive lobular carcinoma (ILC), most of which are oestrogen receptor-positive (ER+). ILCs differ from invasive carcinoma of no special type (IC NST, commonly termed ductal) in three clinically relevant ways: (1) almost universal loss of the cell-adhesion molecule E-cadherin through *CDH1* mutation or silencing (1); (2) a diffusely infiltrative, “single-file” growth pattern that escapes standard radiological assessment and biases trial enrolment; and (3) a distinct metastatic tropism for serosal surfaces and the gastrointestinal tract (2) (3). Despite these features, treatment still follows ductal paradigms—endocrine therapy plus surgery and radiotherapy [1(4)—leaving one-third of patients with endocrine resistance and poorer long-term survival (5).

Progress toward ILC-specific therapy has been hampered by two gaps. First, the RECIST imaging criteria used in most trials under-detect ILC burden, so lobular tumours are often excluded (6). Second, conventional subcutaneous or orthotopic xenografts poorly model ER⁺ disease: engraftment is low and hormone receptor expression is lost (7). Injecting tumour cells directly into the mouse mammary ducts (the MIND model) overcomes these hurdles, yielding ER⁺ and ER⁻ ILC xenografts that recapitulate the full spectrum of progression, dormancy and metastasis (8, 9) (10) (11).

Omics profiling of ILC MIND models and patient tumours has pinpointed an extracellular-matrix (ECM) signature enriched for lysyl-oxidase (LOX) family enzymes that crosslink collagen and stiffen the matrix. Genetic knock-down of LOXL1 or pharmacological blockade with the pan-LOX inhibitor (LOXi) β-aminopropionitrile (BAPN) suppressed tumour growth in proof-of-concept studies (12), but BAPN’s clinical toxicity precludes translation (13).

Recent preclinical studies demonstrate that new LOX inhibitors (LOXis), such as PXS-5505 and LXG6403, enhance chemotherapy efficacy in desmoplastic pancreatic and triple-negative breast cancer models by improving local drug delivery (14–17). PXS-5505 is such a next-generation, highly selective (LOXi) now in phase-I trials for myelofibrosis with a favourable safety profile (18). Here, we repurpose PXS-5505 for ILC using intraductal cell-line and patient-derived xenografts that represent multiple molecular subtypes and disease stages. We demonstrate that LOX inhibition remodels fibrillar collagen, disrupts integrin-mediated signalling and MYC/AP-1 transcriptional programmes, and curtails both primary growth and metastatic dissemination. Collagen fibre metrics and MYC/AP-1 gene signatures emerge as sensitive pharmacodynamic readouts—surrogates that could enable “window-of-opportunity” trials in this imaging-challenged breast cancer subtype.

Collectively, our studies expose a tractable ECM–integrin–MYC vulnerability in E-cadherin-deficient ILC and lay the groundwork for translating PXS-5505, alone or alongside endocrine therapy, into clinical testing for lobular-breast-cancer patients.

## Results

### Quantitative Morphological Features Distinguish ILCs

To capture features that characterise ILCs in an unbiased fashion, we analysed publicly available breast cancer datasets encompassing both transcriptomic and histopathological data (Fig. 1A). We generated a unified transcriptomic landscape by analysing data from 11,000 patient samples from the SCAN-B (19), METABRIC (20), and TCGA (21) cohorts using EMBER (22). As expected, tumours clustered by intrinsic molecular subtypes. While ILCs were predominantly enriched in the Luminal A and Luminal B cluster, a lower proportion were spread across the Basal-like, HER2-enriched and normal-like subtypes (Fig. 1B). This distribution of ILCs across multiple molecular subtypes is consistent with the hypothesis that ILCs may originate from diverse precursor cell populations (23).

**Fig. 1.**
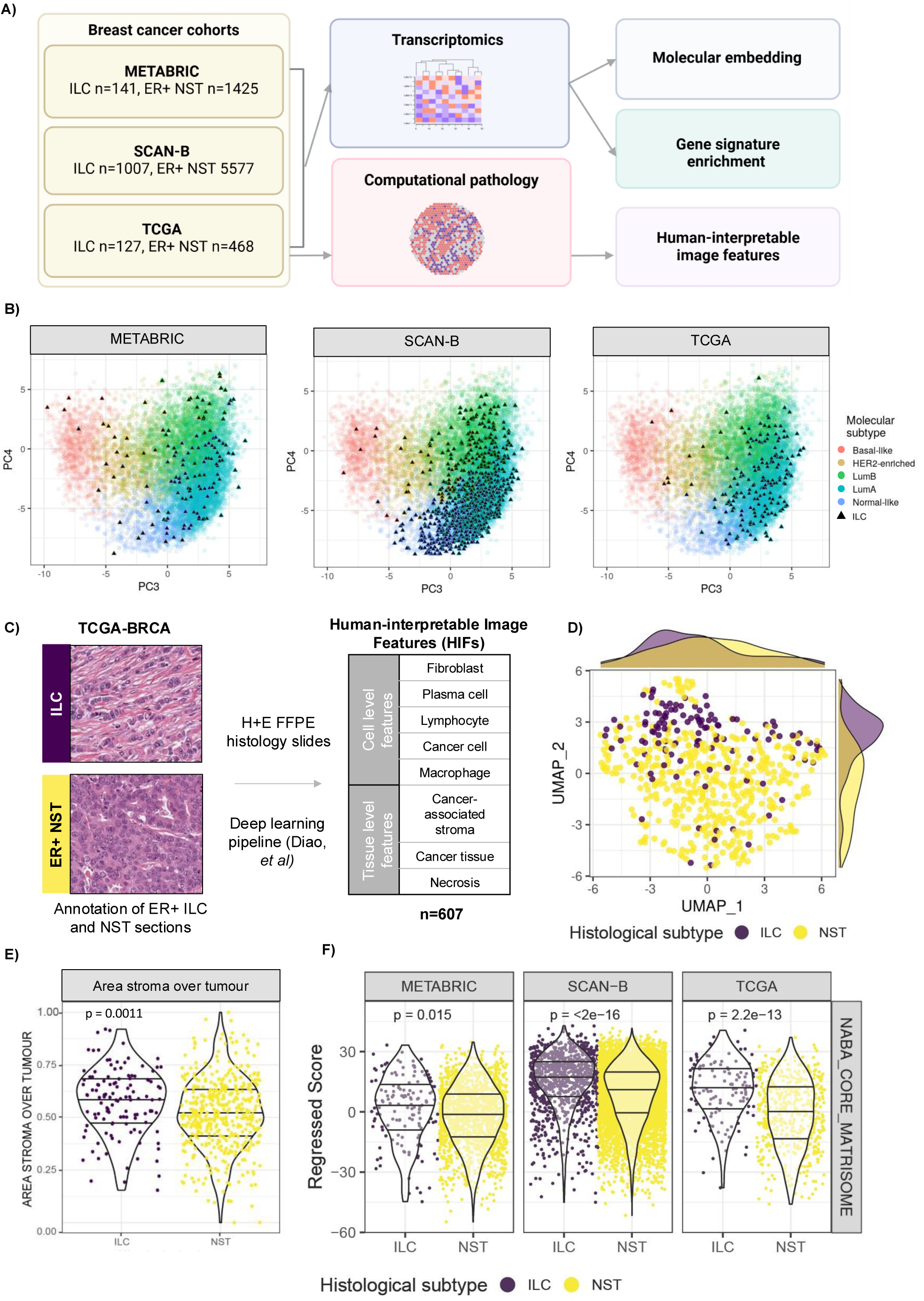
– Quantitative Morphological Features Distinguish ILCs. **(A)** Scheme showing flow of analyses for interrogation of ILC-specific features from breast cancer cohorts. **(B)** EMBER plot of principal components (PC3, PC4) from SCAN-B. ILC samples are denoted by closed triangles. **(C)** Scheme detailing histological feature interpretation derived from (24) **(D)** UMAP of human-interpretable features (HIFs) derived from computational pathology analysis performed in Diao et al. (24) of H+E stained sections from TCGA-BRCA coloured by histological subtype. **(E)** Violin plot showing area of stroma relative to tumour area in H+E stained sections from TCGA ILC and non ILC ER+ (NST) BCs. **(F)** Violin plots showing regressed matrisome signature score in ILC vs non-ILC (ER+ NST) in TCGA, METABRIC and SCAN-B cohorts.

To determine whether morphological features distinguish ILCs from other ER+ BCs in an unbiased manner, we leveraged recent advances in computational pathology. Specifically, we analysed haematoxylin and eosin (H&E)-stained sections from 117 ER+ breast cancer samples in the TCGA BRCA cohort, annotated with 607 cell- and tissue-level features (24) (Fig. 1C). Uniform manifold approximation and projection (UMAP) of these human-interpretable image features (HIFs) revealed a distinct spatial distribution of ILCs compared to NST tumours (Fig. 1D). Features enriched in ILCs included those associated with stromal cell composition, - particularly macrophage density-aligning with previous reports of macrophage enrichment in ILCs (Supp Fig. 1B) (25). Notably, the area of cancer-associated stroma relative to tumour area was significantly greater in ILCs compared to non-ILCs (Fig. 1E).

Given prior evidence that ILCs are enriched for ECM interaction (12, 26), we examined matrisomal gene expression using the core matrisome gene set (NABA_CORE_MATRISOME). Across all three cohorts (SCAN-B, TCGA, and METABRIC), ILCs exhibited higher expression of this core matrisome signature compared to ER+ IC-NSTs (Fig. 1F.) Similarly, the KEGG_ECM_INTERACTIONS and REACTOME_ECM_ORGANISATION gene signatures were enriched (Supp Fig. 1A).

Collectively, these results highlight the intrinsic heterogeneity of ILCs, particularly at the transcriptomic level. The co-occurrence of matrisome gene signature enrichment and stroma-associated morphological features suggest shared biological mechanisms that may represent unique therapeutic vulnerabilities in ILCs.

### Evaluation of the Pan-LOXi, PXS-5505 for the Treatment of MIND Models of ER+ ILC

Prompted by the robust enrichment of ECM-related features in ILC (27), and prior observations that the pan-LOXi, BAPN, reduces ILC xenograft growth (12), we sought clinically translatable strategies to inhibit LOX activity. We selected PXS-5505, a pan-LOXi with demonstrated safety and tolerability in clinical studies, for evaluation in preclinical models of ILC.

To first assess tolerability in immunocompromised *NOD SCID gamma* (*NSG*) females, mice were treated daily with PXS-5505 (10-30 mg/kg daily i.p.) or BAPN (800 mg/kg) for 7 days. BAPN treatment caused significant weight loss (Supp Fig. 2A), while PXS-5505 had no effect on body weight or liver mass (Supp Fig. 2A). A fluorometric activity assay confirmed that PXS-5055 i.p. achieved complete inhibition of the lysyl oxidase family activity in the mouse aorta (Supplementary Figure 2D) (28). Histological evaluation of fibrillar collagen via picrosirius red staining and TWOMBLI analysis (29) showed that PXS-5505 did not alter collagen architecture in the mammary gland, lung, or liver, unlike BAPN which caused changes in collagen fiber morphology (Supp Fig. 2B-C). These findings confirmed that PXS-5505 is well tolerated and does not induce off-target ECM collagen fiber disruption.

**Fig. 2.**
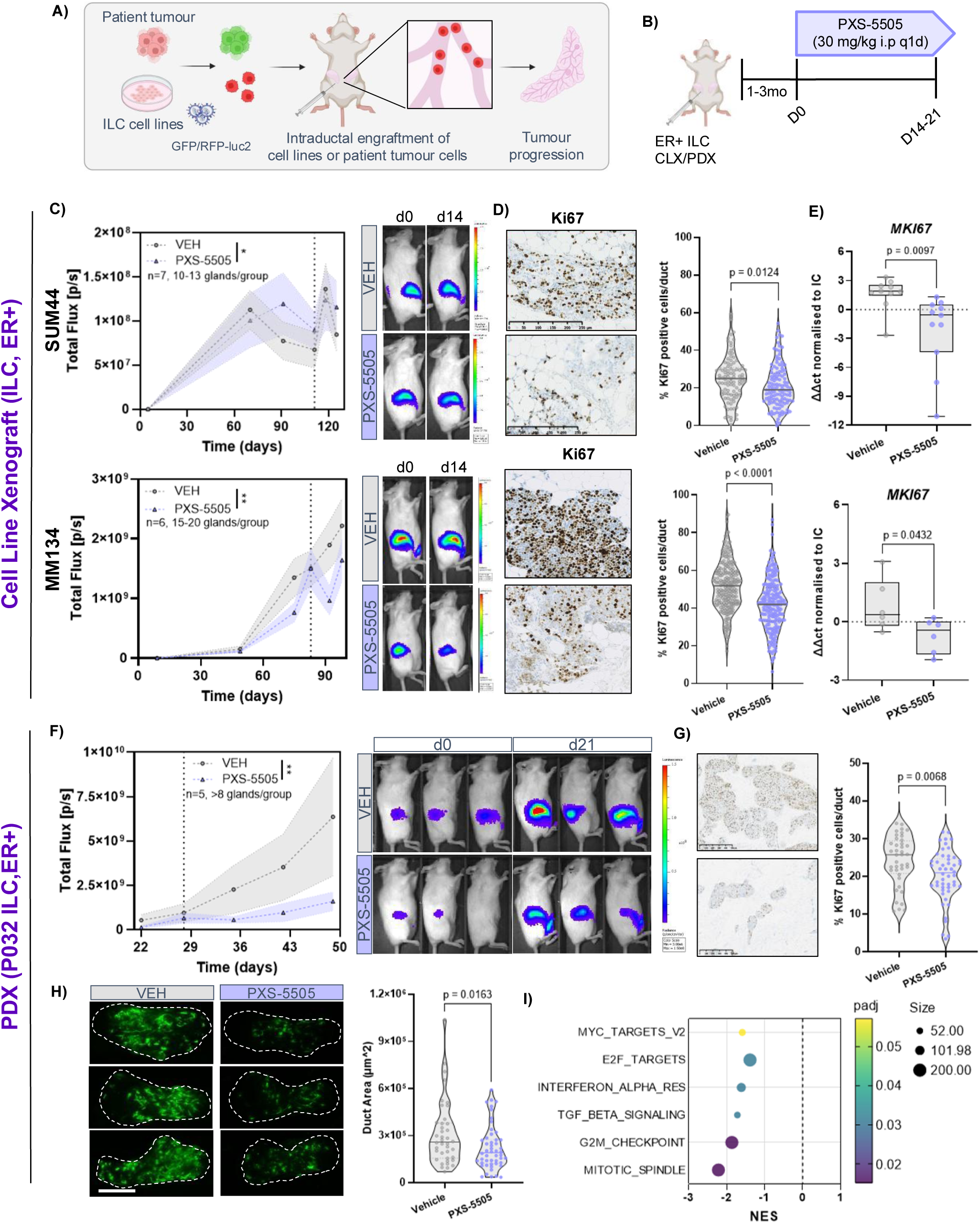
– Evaluation of PXS-5505 for the Treatment of ER+ ILC. **(A)** Scheme for generating cell line (CLX) or patient-derived (PDX) intraductal xenografts. **(B)** Experimental scheme for treatment with 30mg/kg PXS-5505. i.p (intraperitoneal) q1d (1x daily)**(C)** Radiance-based tumour growth measurements SUM44PE-RFP-luc2 (top) and MM134 (bottom) intraductal xenografts. **(D)** % Ki67 positive cells/duct quantified by IHC. **(E)** Expression of MKI67 quantified by qRT-PCR. **(E)** Radiance-based tumour growth measurements of an ER+ ILC intraductal PDX transduced with GFP-Luc2 and treated with PXS-5505. Representative bioluminescence images shown (n=5/ group). **(G)** % Ki67 IHC positive cells/duct quantified by QuPath **(H)** Representative fluorescence stereomicrographs of engrafted inguinal glands. Area of tumour-cell filled ducts, n=5 mice, 8 glands/groups. **(I)** Normalised enrichment scores (NES) for HALLMARK gene sets in PXS-5505 treated vs Vehicle tumours (n=4/group). Significance determined by Students t-test or ANOVA (Tukey’s multiple comparisons) and expressed as * P < 0.05, ** P < 0.01, *** P < 0.001. Analysis of bioluminescence growth curves carried out using linear splines to fit the growth curves and the lr-test, using maximum likelihood estimation, to check for differences. Data are shown as mean of 3 or more biological replicates ± SEM.

Next, we evaluated the efficacy of PXS-5505 using MIND xenograft models of ER+ ILC. We intraductally engrafted two RFP-Luc2 tagged ER+ ILC cell lines, SUM44PE and MDA-MB-134IV (MM134), into 2-4 mammary glands of NSG females (Fig. 2A). After three months, when radiance reached 10^7^-10^8^photons/sec/gland, mice were treated with PXS-5505 (30mg/kg/d) i.p. for 14 days (Fig. 2B). Despite the indolent growth of these ILC models, PXS-5505 significantly reduced the slope of tumour growth curves, with a stronger effect observed in the faster-growing MM134 model (p<0.01) than in the SUM44 model (p<0.05) (Fig. 2C).

At study endpoint, engrafted glands were harvested and cell proliferation assessed. LOXI treatment reduced Ki67 expression in both models compared to vehicle, with Ki67 indices reduced by -16.1% in SUM44 (p<0.05), and -18.7% inMM134 xenografts (p<0.01) (Fig. 2D). Concordantly, *MKI67* mRNA levels were reduced 1.13-fold in SUM44 and 1.42-fold in MM134 (Fig. 2E). Thus, PXS-5505 has antiproliferative effects and Ki67 expression may serve as a more sensitive biomarker of drug response than the radiance measurements in these slow-growing models.

Next, we sought to test the drug in a PDX generated from a pleural effusion derived from of a heavily pre-treated patient diagnosed with ER+ PR+ HER2-ILC (P032). PDXs from ILCs are challenging to establish and often require up to one year before the engrafted cells have grown to sufficient numbers to allow for serial passage (10, 30, 31). At generation 5 (G5), the grafts retained ER+, PR+, HER2-status as confirmed by IHC. One month post-engraftment, once luminescence reached 10^7^ photons/sec/gland, mice were treated with PXS-5505 for 3 weeks. Treatment resulted in reduced radiance (Fig. 2F) and fluorescence stereoscopy at endpoint revealed diminished signal intensity and extent in treated glands(Fig. 2H). Quantification confirmed significant reductions in ductal area (Fig. 2H) and % tumour cellularity (Supp. Fig. 3A) upon treatment. IHC revealed a decrease in Ki67 index of -16.5% (Fig. 2G). Transcriptomic profiling of the engrafted glands showed significant downregulation of cell proliferation-associated pathways, including G2/M checkpoint and mitotic spindle, as well as IFN-α response and TGF-β signalling, following PXS-5505 treatment (Fig. 2I).

Collectively, these results demonstrate that PXS-5505 inhibits primary tumour growth in multiple preclinical ILC models. Histological analyses and Ki67-based assessments appeared more sensitive than radiance imaging for detecting therapeutic effects in indolent ILC models.

### LOXi inhibits ILC Progression in Window-of-Opportunity Trial Settings

To further clinical translation of our findings, we designed experimental paradigms that emulate the structure of window-of-opportunity trials (WoO). These trials are used to introduce new drugs into clinics, typically involve short-term administration of a therapeutic agent -often for up to four weeks-between diagnostic biopsy and surgical resection. The predictive value of WoO was clinically validated in ER+ breast cancer patients (32, 33).

We first assessed the effect of ER degradation on ILC cell proliferation *in vitro*. SUM44 cells, which proliferate more slowly than MM134 cells, were more sensitive to the selective ER degrader (SERD) fulvestrant and showed a 36.2% decrease compared to control at endpoint. MM134 cells showed a 20.8% decrease and as expected, the ER-IPH-926 model was unresponsive to fulvestrant (Fig. 3A).

**Fig. 3.**
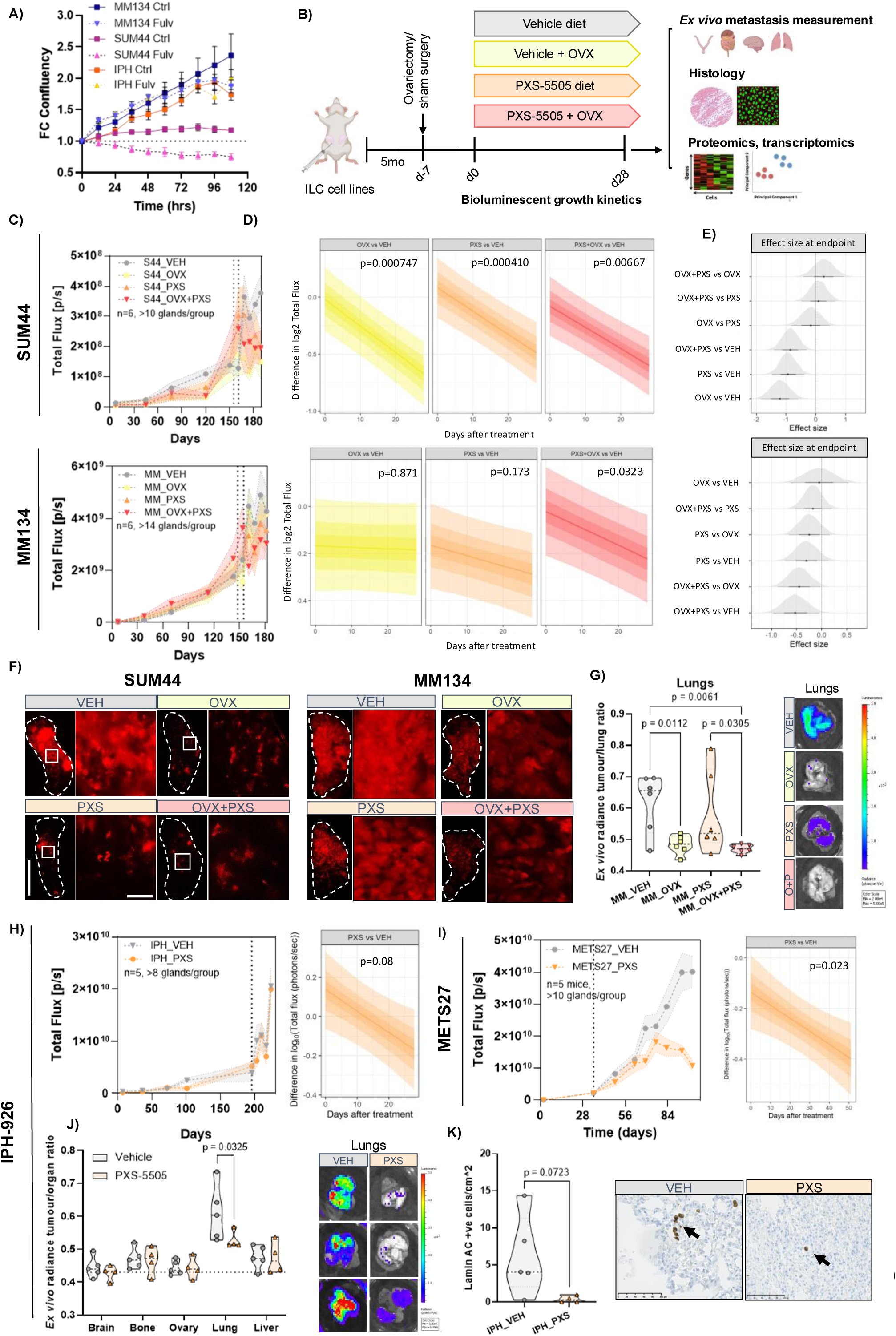
-LOXi inhibits ILC Progression in Window-of-Opportunity Trial Settings. **(A)** Relative confluency of MM134 and SUM44 cells over time treated with selective ER degrader (SERD) Fulvestrant (Fulv). **(B)** Scheme of in vivo experiment **(C-E)** Radiance of SUM44-RFP-Luc2 (top) and MM134-RFP-Luc2 (bottom) intraductal xenografts (n=5-6/group). Dashed line indicates surgery and start of treatment. **(F)** Representative fluorescence stereomicrographs of engrafted glands at endpoint. Dotted line highlights circumference of engrafted inguinal mammary gland. Scale bars 1cm (left) and 2mm (right) **(G)** Ex vivo radiance of lungs from NSG hosts bearing intraductal MM134 xenografts at endpoint. Values normalised to tumour burden. Representative stereomicrographs of lungs overlaid with scaled bioluminescence. **(H-I)** Tumour xenografts radiance of IPH-926 and METS27 (PDX) xenograft bearing mice**. (J)** metastatic organs at endpoint**. (K)** Bar graph/ photomicrographs of lung sections stained with anti-human specific Lamin AC antibody counterstained with haematoxylin. Arrows indicate disseminated human tumour cells. Significance determined by students or multiple t-tests and expressed as * P < 0.05, ** P < 0.01, *** P < 0.001. Data are shown as mean of n>10 glands ± SEM.

To mimic clinical treatment timelines, we extended the duration of PXS-5505 administration in xenografted mice to four weeks. To facilitate this and to avoid repeated i.p. injections, PXS-5505 was delivered through a custom-formulated diet. Because aromatase inhibitors (AIs) – the standard of care for postmenopausal patients with ILC- are ineffective in mice due to species-specific differences in aromatase biology (34), we employed ovariectomy (OVX) to deplete endogenous 17-β-estradiol (E2) and mimic AI effects (Fig. 3B). For the TN IPH-926 model, PXS-5505 was tested as a monotherapy.

At 5-7 months post engraftment, when luminescence reached 10^8^ photons/sec/gland in SUM44 and 10^9^ in MM134 xenografts, mice underwent either OVX or sham surgery. Seven days later -allowing sufficient time for depletion of circulating ovarian hormones-PXS-5505 was introduced in half of the OVX and control mice. Tumour progression was monitored weekly via in vivo bioluminescence imaging, and endpoint analyses were conducted on blood and tissues (Fig. 3B). LOX activity assays on the mouse aortas confirmed on-target drug activity at endpoint with a significant reduction in enzymatic activity (Supp Fig. 2D). While OVX led to the expected increase in body weight, dietary PXS-5505 had no effect on weight nor general condition, confirming good tolerability (Supp Fig. 2E).

In agreement with in vitro findings (Fig. 3A), OVX significantly reduced radiance in SUM44 (p<0.001) but not in MM134 xenografts (Fig. 3C, D). To analyse treatment-specific differences in growth rates, we used BioGrowler a Bayesian hierarchical modelling tool for longitudinal data analysis (35). In SUM44 xenografts, PXS-5505 monotherapy significantly reduced tumour growth (p<0.001) with a similar magnitude as OVX or the combined treatment (p<0.01) (Fig. 3D).

Effect size estimates based on the slope differences of growth curves were similar for OVX, PXS-5505, and their combination (effect sizes -0.8 to -1.2) (Fig. 3E). In MM134 xenografts, OVX alone had no effect, but PXS-5505 monotherapy showed a trend towards growth inhibition (p=0.17). Combined OVX+PXS-5505 treatment significantly reduced tumour growth compared to both vehicle (p=0.032) and OVX alone (p=0.045) (Fig. 3D, E). Histological analysis of H+E-stained sections showed characteristic diffuse growth in the SUM44 model, and a reduction in tumour cellularity from 6.5% in the vehicle to 5%, 3.3% and 4.6% in the OVX, PXS and O+P conditions (Supp Fig. 3B). The MM134 model showed extensive tumour infiltration in vehicle-treated mice covering 54.1% of total gland area. This was reduced to 27.3%, 47.4% and 38.1% in the OVX, PXS-5505 and combination groups, respectively (Supp Fig. 3C).

In the slow growing SUM44 model, *ex vivo* radiance imaging of distant organs showed no detectable metastatic dissemination at this stage. In contrast, in mice bearing MM134 xenografts lung radiance was increased at endpoint (Fig. 3G). OVX alone reduced this metastatic burden (p=0.01), with a further decrease observed in the OVX+PXS-5505 combined group (p=0.006) (Fig. 3G).

To test PXS-5505 in a TN ILC model, we used IPH-926RFP-Luc2 cells, derived from the ascitic fluid of a pre-treated patient (36, 37). Seven months post-engraftment, when radiance reached 10E9/sec/gland, treatment was initiated. The effect of PXS-5505 failed to reach significance over 30 days (p=0.08) (Fig. 3H). However, *ex vivo* imaging revealed reduced lung radiance in treated mice (Fig. 3J) and quantification of IHC of lung sections for Lamin A/C demonstrated a decrease in the number of tumour cells per cm^2^ of lung tissue of treated animals indicating reduced metastatic load (Fig. 3K).

From a pleural effusion (METS27) of a patient with ER+ ILC who relapsed one-year post-diagnosis and paclitaxel treatment, we successfully passaged and expanded cells. At generation 4, we were able to test two different conditions and compared PXS-5505 versus vehicle. Sixty days of treatment significantly reduced in vivo luminescence (p=0.023) (Fig. 3I). In summary, across three cell line-derived xenografts and 2 PDX models representing different disease stages and molecular subtypes, LOXi -alone or combined with hormonal deprivation-reduced primary ILC growth and/or metastasis. These findings underscore the therapeutic potential of this drug in inhibiting ILC progression.

### Exploring Alternative Histological Endpoints in ILC

ILC disease burden assessed by imaging often fails to match that revealed by histopathology (38, 39). Furthermore the utility of the Ki67 index, often used as gold standard pharmacodynamic endpoint in window-of-opportunity trials, is debated because it is low in these slow growing tumours (40). In the SUM44 model, the Ki67 index significantly declined from 21.9% in vehicle-treated controls to 14%, 14.8% and 11.4% in OVX, PXS-5505 and combination groups, respectively (p<0.0001 for all comparisons; Fig. 4A, top), in concordance with the radiance-based tumour growth measurements. Interestingly, in the MM134 model -which did not show radiance reduction with OVX - Ki67 index nonetheless dropped from 35.4% to 14.8% upon OVX alone (p<0.0001). PXS-5505 monotherapy reduced Ki67 to 27.3% (p=0.0002), while the combination OVX+PXS-5505 yielded the most substantial reduction, to 10.2% (Fig. 4A, bottom).

**Fig. 4.**
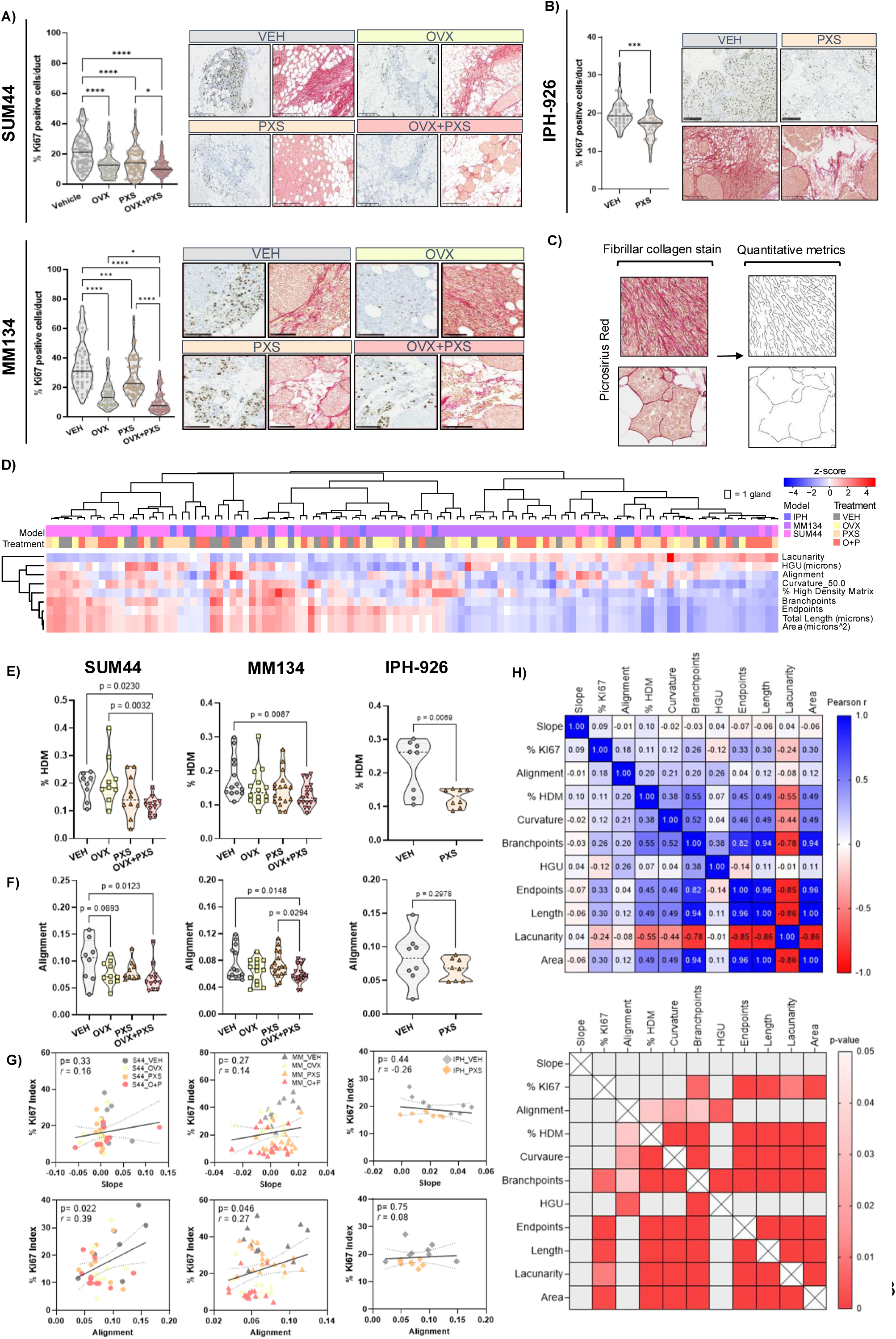
-Exploring Alternative Histological Endpoints (A-B) Violin plot showing % Ki67 positive tumour cells/duct quantified by IHC in SUM44 (top), MM134 (bottom) and IPH-962 (right) MIND xenografts. Representative micrographs of anti-Ki67 antibody stained and PSR stained histological sections (20X magnification, scale bar = 100µm). **(C)** Collagen feature identification and quantification analysis pipeline of PSR stained sections using TWOMBLI. **(D)** Heatmap of collagen morphometrics in treated and untreated SUM44, MM134 and IPH-926 tumours. **(E-F)** Violin plots showing collagen fiber density (% high density matrix) and alignment in treated and untreated xenograft models. **(G)** Correlation plots showing Ki67 index vs radiance (left) and collagen fiber alignment (right). Pearson correlation r and p-values indicated. **(H)** Pearsons correlation matrices for imaging, proliferation and collagen metrics from SUM44, MM134 and IPH-926 models combined (top), and corresponding p-value for each comparison (bottom). Significance determined by one-way ANOVA and expressed as * P < 0.05, ** P < 0.01, *** P < 0.001. Data are shown as mean of 5 or more individual glands.

The reductions in KI67 index were similar in both the IPH-926 model (Fig. 4B) and the METS27 model (Supp Fig. 3D,E) (-16.5% and -11.39% respectively), despite PXS-5505 treatment significantly reducing tumour radiance in the METS27 model but not the IPH-926 model. This further reflects the discordance between Ki67 index and tumour burden and suggests that alternative endpoint measures should be considered.

To assess LOXi effects more directly, we analysed fibrillar collagen architecture revealed by picrosirius red staining of xenografted tissue. Quantitative analysis using TWOMBLI (29) to measure fiber length, alignment, density, and branching (Fig. 4C) revealed substantial intra-and inter-model heterogeneity, mirroring the histopathologic variability seen in human ILC specimens (Supp. Fig. 3B, 4A) (41). Hierarchical clustering indicated that LOXi-treated glands were enriched in clusters characterised by reduced fiber length, area, and branching (Fig. 4D). Importantly, both collagen matrix density (% HDM) (Fig. 4E) and fiber alignment (Fig. 4F) were significantly decreased in the combination group compared to vehicle in SUM44, and MM134 and by LOXi monotherapy in IPH-926 xenografts (Fig. 4E,F).

To examine the relationship between tumour matrix remodelling, cell proliferation, and tumour growth kinetics, we constructed Pearson correlation matrices linking imaging, Ki67 index, and different collagen metrics. Linear regression analysis confirmed a lack of correlation between the luminescence-based growth rate (slope) and Ki67 index (Fig. 4G, top). It revealed a positive correlation between fiber alignment and Ki67 index in SUM44 (n=39, r=0.37, p=0.029) and MM134 (n=59, r=0.26, p=0.046) models (Fig. 4G, bottom) but not in the TN IPH-926 model where the magnitude of response to LOXi by Ki67 was less(n=16, r=0.08 p=0.75). Across all models, the Ki67 index correlated with fiber branch points, area, and length (Fig. 4H). When the TN IPH-926 model was excluded from the analysis (given its reduced treatment responsiveness), additional significant correlations emerged between Ki67 and fiber alignment and curvature.

These findings suggest that combining alterations in collagen architecture and Ki67 indices may offer superior sensitivity of LOXi response when evaluating therapeutic efficacy as surrogate markers possibly enabling future clinical trials targeting the lobular BC subtype.

### The ILC-Intrinsic Matrisome Is Modulated by LOXi

To investigate how LOXi-induced collagen fiber remodelling impacts the ECM and subsequently influences ILC cell behaviour, we performed tandem mass tag (TMT) proteomics on xenograft samples treated for 28 days with LOXi. Proteins were identified by aligning peptide sequences to both human and murine reference databases, allowing for discrimination between tumour (human) and host (murine)-derived proteins. Consistent with the slower and more diffuse growth pattern of the SUM44 model, fewer proteins were detected compared to the more proliferative MM134 xenografts (8043 vs 11161). The percentage of human proteins identified was 46.9% in the SUM44 model and 61.40% in MM134.

In SUM44 xenografts, OVX significantly altered the expression of 1376 human proteins (p<0.05, |log2(FC)| >0.5), with 90% of these downregulated (Fig. 5A). PXS-5505 monotherapy resulted in 583 differentially expressed (DE) proteins (Fig. 5B). In the OVX+PXS group only DE 325 proteins were found, likely due to the diminished number of tumour cells present following the effective treatment, as suggested by radiance and histological analyses (see Fig. 3F, Supp Fig. 3B). In MM134 xenografts, OVX alone and OVX+PXS-5505 treatment modulated a similar number of proteins-659 and 640 and PXS-5505 monotherapy regulated 234 proteins, most DE proteins being downregulated (Fig. 5B, Supp Fig. 5A).

**Fig. 5.**
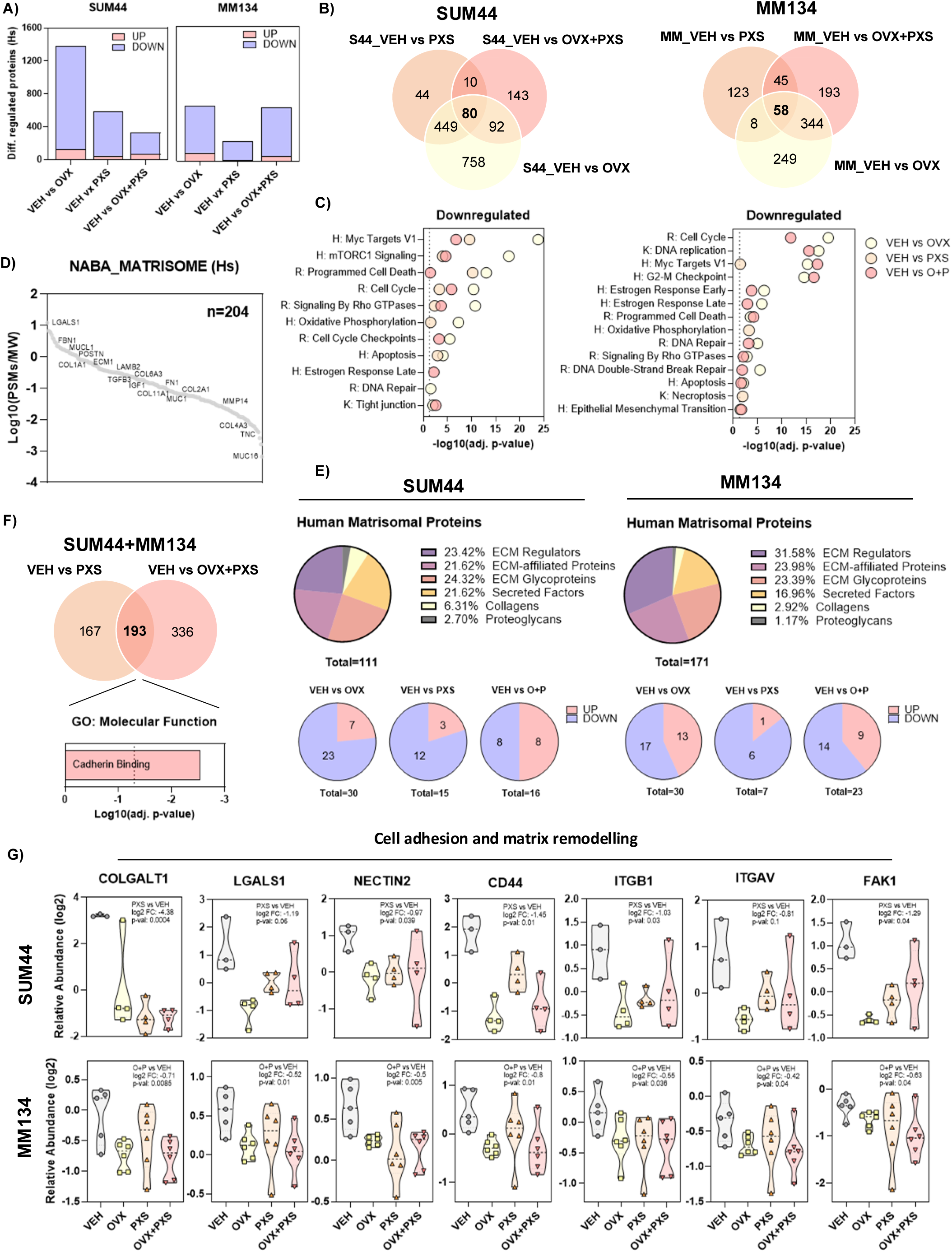
– The ILC-Intrinsic Matrisome Is Modulated by LOXi. **(A)** Bar plots showing Multiplex proteomic analysis of SUM44 and MM134 xenografts. Number of detected proteins for each experimental condition indicated (top) and differentially regulated proteins identified for each comparison. Threshold p-value<0.05, proteins with log2(FC) >0.5 in red and log2FC <0.5 in blue. **(B)** Venn diagram of overlapping differentially expressed (DE) proteins in SUM44 (left) and MM134 (right). **(C)** Pathway enrichment (HALLMARK, REACTOME, KEGG) for downregulated proteins in each condition vs Vehicle. **(D)** Human matrisome protein abundance in ILC xenografts as ranked by log10 peptide specific matches (PSMs)/ molecular weight. **(E)** Pie charts of human matrisomal proteins identified by MatrisomeAnalyzeR (top). Pie charts of differentially regulated matrisomal proteins in each treatment condition (bottom). **(F)** Venn diagram of commonly regulated proteins in VEH vs PXS and VEH vs OVX+PXS conditions in both models. Functional enrichment by STRING of common differentially regulated proteins. **(G)** Abundance of matrix and cell adhesion related proteins significantly differentially regulated by PXS in SUM44 (top) and by OVX+PXS in MM134 (bottom). Statistical significance determined by one-way ANOVA and expressed as * P < 0.05, ** P < 0.01, *** P < 0.001.

Pathway enrichment analysis of the downregulated proteins performed using the KEGG, REACTOME, and HALLMARK gene sets-revealed broad suppression of ER signalling, cell cycle progression and apoptosis-related pathways in response to OVX in both models as expected (Supp Fig. 5B), in light of the reduction in cellular proliferation observed via Ki67 staining (see Figure 4A). Notably, all three treatment conditions in the SUM44 model led to suppression of cell cycle-related pathways, whereas in MM134, significant suppression was observed only in the OVX and OVX+PXS-5505 groups (Supp Fig. 5B). Protein-level reductions in key markers of proliferation (e.g., *MKI67, PCNA, MCM6*) and the cell cycle regulation (e.g., *CDK1*) further support this observation (Supp Fig. 5C). Interestingly, across both models the only pathways consistently modulated by LOXi monotherapy were MYC Targets V1 and Oxidative Phosphorylation. This suggests a more limited metabolic and transcriptional effect of LOX blockade in the absence of systemic E2 depletion (Fig. 5C).

To further dissect the composition of the ILC cell-derived matrisome and its modulation by LOXi, we annotated human ECM components using *MatrisomeAnalyzeR* (42). Ranking of human matrisome proteins averaged across the models identified Galectin-1 (*LGALS1*) as the most abundant component (Fig. 5D). Galectin-1 is an immunomodulatory protein associated with high-grade breast tumours (43), known to promote fibrosis (44) and to regulate ECM remodelling, cell adhesion, and invasion through interactions with ECM components. While ECM deposition is typically attributed to cancer-associated fibroblasts (45), our proteomic analysis - based on human-specific peptide sequences-conclusively demonstrates ILC cells synthesise and remodel ECM proteins. Some components were model-specific; for instance, *COL1A1* was detected exclusively in the SUM44 model, whereas *COL18A1* was unique to the MM134. Others, including *COL6A3*, Fibrillin-1 (*FBN1*) and *MMP14*, were present in both (Supp Table 1). The overall proportion of these proteins and the distribution across matrisome categories - collagens, laminins, glycoproteins, and ECM regulators-was comparable between models (Fig. 5E). Across all 3 treatment conditions matrisomal proteins were among those differentially regulated, with the greater proportion downregulated (50-85%) (Fig. 5E) suggesting a robust suppression of ECM production and remodelling activity in response to LOXi.

Protein-protein interaction enrichment analysis was carried out using STRING (46). Network enrichment of proteins significantly modulated by PXS and OVX+PXS (n=193) revealed cadherin binding as a top enriched functional category (Fig. 5F) suggesting alterations in cell-cell and cell-ECM adhesion dynamics. Several key matrix sensing and mechano-transduction proteins were downregulated, including *CD44* – a receptor for ECM ligands that can regulate cell adhesion and survival, *ITGB1* - key collagen binding integrin and FAK (*PTK2*), a central downstream mediator thereof (Fig. 5G). Thus, LOXi treatment not only disrupts collagen fiber architecture and density, but results, directly or indirectly, in downmodulation of adhesion receptors and associated signalling proteins. This suggests a dual mechanism whereby LOXI affects both the structural ECM and the cellular machinery that interprets matrix-derived cues.

### Dependency on ITGAV/ITGB5 is a Targetable Vulnerability in *CDH1* Deficient Tumour Cells

To impair ILC cell fitness, LOXi-induced fiber and matrix changes must, at some level, target a tumour-cell intrinsic dependency. Given this, we searched for ILC cell intrinsic dependencies that could explain the observed sensitivity to LOXi. Given E-cadherin loss is a common feature of ILCs, we analysed genome-wide CRISPR-Cas9 screen data from the DepMap initiative (47) to identify genes selectively essential in E-cadherin defective cells (i.e. E-cadherin synthetic lethal genes (48, 49)), reasoning that these would be good candidates for ILC tumour cell intrinsic dependencies. (Fig. 6A). We classified 1673 tumour cell lines in DepMap by E-cadherin (*CDH1*) mRNA expression into E-cadherin high (>2) and low (<2) groups (Fig 6B), and interrogated genome-wide CRISPR-Cas9 screen data to find gene more essential in the low (i.e. candidate E-cadherin synthetic lethal effects, illustrated by negative effect sizes in Fig 6C). This identified genes such as *CHMP4B* previously shown to be essential in tumour cells with deletion of *CDH1* on chromosome 16q (Fig 6C) (50), validating our approach. Analysis also revealed E-cadherin synthetic lethal effects associated with two genes, *ITGAV* and *ITGB5*, that encode the heterodimeric integrin receptors, which sense the biomechanical properties of collagens (51) (Fig. 6C). *ITGAV* encodes the alpha chain V, whereas *ITGB5* encodes integrin beta-5 which interacts with and signals via FAK, Annexin V and PAK4. A similar analysis to that described in (52), where tumour cell lines were not classified by E-cadherin expression but defined by their epithelial or mesenchymal status also identified *ITGAV* and *ITGB5* as selective dependencies in mesenchymal cells (Fig. 6D). This suggests that *ITGAV* and *ITGB5* functions are essential in tumour cells that have lost E-cadherin expression, but also in those that have undergone EMT via other routes (Fig. 6E, F). Mass spectrometry-based proteomic data from 52 ER-positive breast cancers in the TCGA study—including 6 tumours with deleterious truncating mutations in CDH1—were analysed. Normalized protein expression levels of ITGAV and ITGB5 were higher in CDH1-mutant (E-cadherin deficient) tumours compared to wild-type, though the differences were not statistically significant (p=0.08 and p=0.13, respectively) (Supp Fig. 6A).

**Figure 6.**
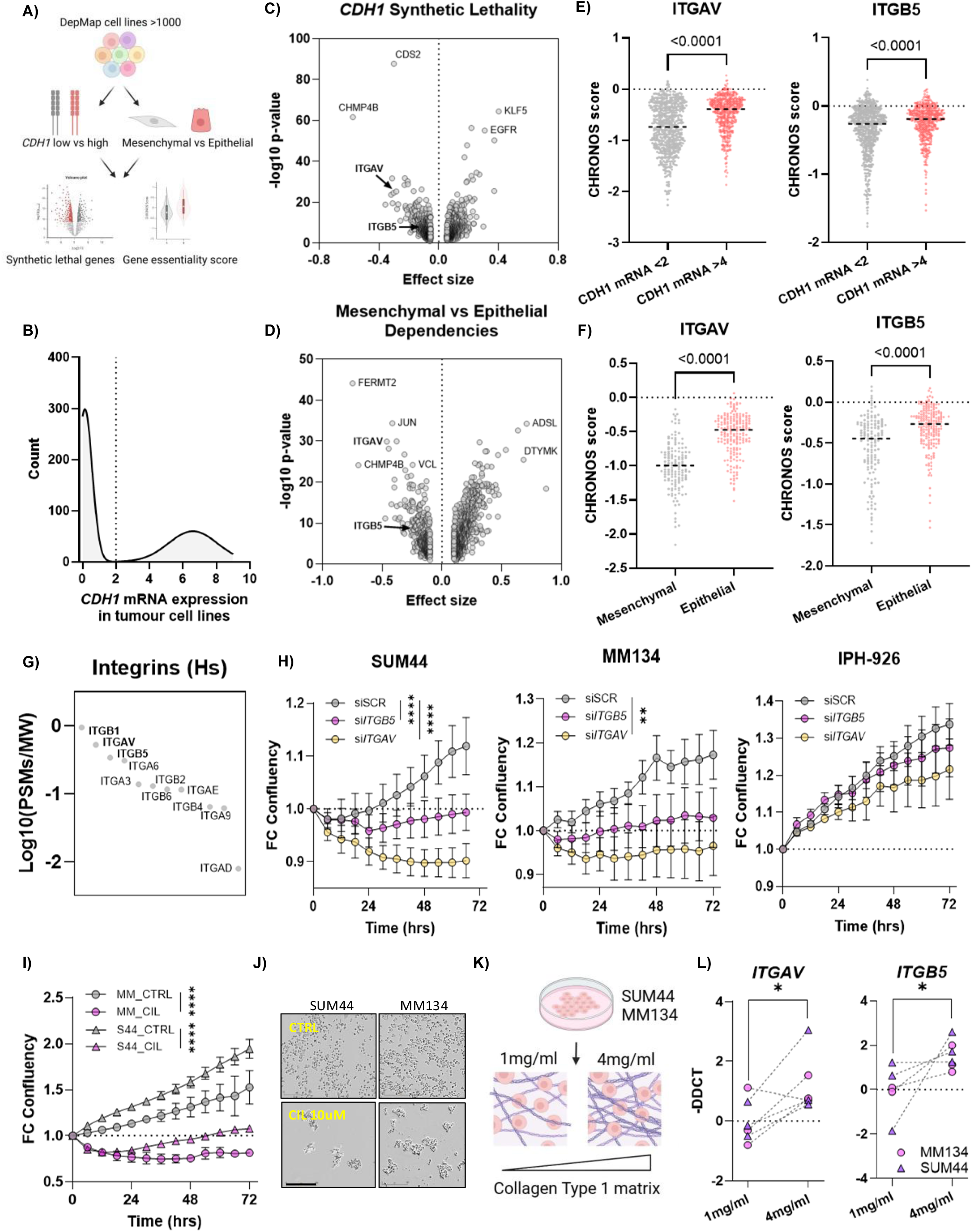
-. **(A)** Schematic of analyses carried out to identify E-cadherin synthetic lethal effects or essential genes in mesenchymal tumour cell lines. Tumour cell lines described in the DepMap dataset (47) were classified in two ways: (i) as either having low or high E-cadherin (CDH1) mRNA expression (left); or (ii) as having a mesenchymal or epithelial morphology, based on a pre-defined (47). **(B)** Histogram illustrating E-cadherin (CDH1) mRNA expression in 1673 tumour cell lines. The threshold of normalised expression of 2 was used to dichotomise the tumour cell line panel into E-cadherin high (mRNA normalised expression > 2) and low (mRNA normalised expression < 2) cohorts. **(C)** Volcano plot illustrating gene dependency analysis based on E-cadherin mRNA classification described in (B). Negative Effect Size effects reflect candidate E-cadherin synthetic lethal genes. **(D)** Volcano plot illustrating gene dependency analysis based on mesenchymal vs. epithelial classifications. Negative Effect Size effects reflect candidate dependencies in mesenchymal tumour cell lines. **(E, F)** Scatter plots illustrating ITGAV gene dependency/synthetic lethality in E-cadherin low (CDH1 mRNA <2) vs. high low (CDH1 mRNA >2) tumour cell lines (shown in E) and in mesenchymal vs. epithelial tumour cell lines (shown in F). Each data point represents a single tumour cell line. Negative CHRONOS scores depict the scale of gene dependency. P values calculated by Mann-Whitney test. **(G)** Abundance of integrins in ILC xenografts. **(H)** Fold change confluency of ILC cell lines transfected with siRNA directed against ITGAV and ITGB5 as well as a scrambled control (siSCR). **(I-J)** Fold change confluency of cells treated with 10µM cilengitide (CIL). Representative images shown. **(K)** Culture of ILC cell lines in 3D collagen type 1 matrices of increasing density. **(L)** Expression of ITGAV and ITGB5 in ILC cells in 1mg/ml vs 4mg/ml collagen matrices determined by qRT-PCR. Statistical significance determined by two-way ANOVA or students t-test and expressed as * P < 0.05, ** P < 0.01, *** P < 0.001.

In parallel, our proteomics analysis of ILC models identified *ITGB1, ITGAV*, and its heterodimer partner *ITGB5* as the three most abundant integrins in the two ILC models above (Fig. 6G). In line with the hypothesis that LOXi interferes with critical interactions of ILC cells with collagen fibers, treatment with LOXi not only disrupted collagen fibers but also led to the downregulation of *ITGB1* and *ITGAV* protein expression in both models (Fig. 5H) likely by an indirect mechanism. the disruption of collagen fibres result of inhibition of LOX has been shown to reduce fibronectin assembly (16, 53), hence less of the RGD-peptide on fibronectin which is a ligand of integrin αvβ5 heterodimers is available with downregulation of the receptor as a consequence.

Functional dependence of *CDH1* deficient ILC cell lines on the integrins identified in the genetic screens was confirmed through knockdown of *ITGAV* and *ITGB5* using siRNA (Supp 6B-C). Silencing of *ITGAV* decreased survival in all three models with 25% and 18% over 72hrs in SUM44 and MM134 cells, and 9% in the IPH-926 cells. Silencing of *ITGB5* affected cell proliferation less reaching significance only in the SUM44 model (Fig. 6H). Pharmacological inhibition with cilengitide (CIL), a selective αvβ3/αvβ5 inhibitor, induced rapid cell detachment in 2D cultures (Fig. 6I-J) further confirming αvβ3/αvβ5 as a functional dependency of ILC cells. To explore the influence of collagen density on integrin expression, ILC cells were cultured in collagen type 1 matrices of concentrations reflecting physiologic (1mg/ml) versus tumour-like (4mg/ml) ECM stiffness (54) (Fig. 6K). Culture in high-density collagen matrices increased mRNA expression of both *ITGAV* and *ITGB5* transcripts (Fig. 6L).

The relatively low transcript expression of *ITGB1, ITGAV* and *ITGB5* in the ILC cell lines SUM44 and MM134 compared to BC cell lines in the Cancer Cell Line Encyclopaedia (CCLE) (Supp Fig. 6D), is consistent with previous findings reporting breast cancer cells with low integrin expression have increased sensitivity to integrin modulation (55). Together, these findings support a model in which loss of E-cadherin renders ILC cells reliant on the ECM-engaging integrins ITGAV and ITGB5.

The down modulation of MYC target genes in our proteomic analysis (Fig. 5D) pointed to reduced MYC activity, a potential downstream consequence of integrin signalling disruption (56, 57). Transcription factor interaction mapping using PathwayNET (58) identified MYC, JUN, FOS and NFKB1 at the centre of a network regulating key ECM and adhesion-related genes (Supp. Fig. 6F). Treatment of ILC cells embedded in high collagen matrices with cilengitide led to a reduction in *MYC* transcript levels, suggesting MYC activity may be reciprocally maintained by integrin signalling under matrix dense conditions (Supp. Fig. 6E) (59).

### LOXi Supresses ILC-Specific Transcriptional Programs Driven by ECM-Dependent Signalling

To delineate the transcriptional networks modulated by LOXi in ILC models, we first analysed proteins downregulated by LOXi or OVX using over-representation analysis testing for the enrichment of transcription factors identified by ENCODE ChIPseq dataset (60). In the SUM44 model, gene products downregulated across all treatment conditions, showed significant enrichment for MYC transcription factor targets, implicating reduced MYC activity in mediating treatment-induced gene expression changes. In the MM134 model this enrichment was also evident with LOXi monotherapy but more pronounced in the combination group (Supp Fig. 5B).

To further assess the transcriptional effects of LOX inhibition, we performed RNA sequencing on xenografted glands treated with LOXi. As expected, Gene set enrichment analysis (GSEA) showed that OVX suppressed estrogen response and cell cycle-associated pathways (e.g. G2M Checkpoint, E2F Targets) (Fig. A). TNFα Signalling via NFκβ and MYC-related pathways were most significantly downregulated by LOXi in both models (Fig. 7A).

**Figure 7.**
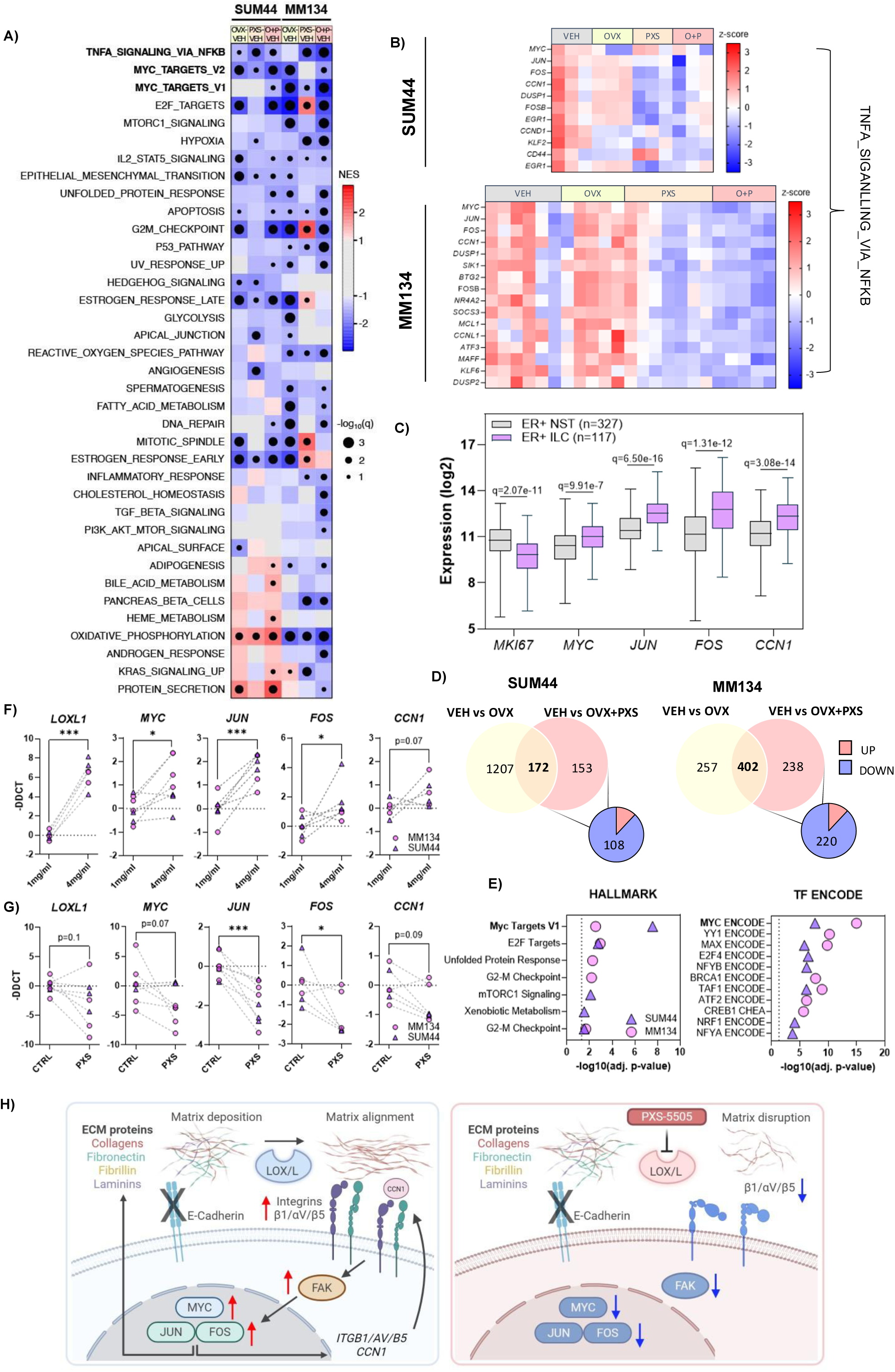
-L**O**Xi **Supresses ILC-Specific Transcriptional Programs (A)** Heatmap showing normalised expression score (NES) values for HALLMARK gene sets from RNAseq profiling across all conditions vs Vehicle. Only the HALLMARK gene sets passing significance (abs(NES) > 1, q-value < 0.1), in at least one condition, are shown. HALLMARK gene sets are ordered from lowest mean NES across conditions (top) to highest NES across conditions (bottom). **(B)** Expression (z-score) of genes contributing to enrichment for TNFA_SIGNALLING_VIA_NFKB and differentially regulated by PXS alone or OVX+PXS. **(C)** Expression of MYC, JUN, FOS and CCN1 in ER+ ILC vs ER+ NST’s from TCGA cohort. **(D)** Venn diagram of commonly regulated proteins in Veh vs OVX and Veh vs OVX+PXS (orange), and only differentially regulated in the Veh vs OVX+PXS condition (grey). Direction of regulation indicated in pie chart. **(E)** HALLMARK pathway enrichment (left) and TF target enrichment (right) for proteins exclusively regulated by OVX+PXS. TF motifs were obtained from ChEA2016 + ENCODE collected in the EnrichR database. **(F)** ILC cells were cultured in collagen matrices. Expression of LOXL1, MYC, JUN, FOS and CCN1 determined by qRT-PCR in 1mg/ml vs 4mg/ml collagen matrices. **(G)** ILC cells cultured in 3D collagen matrices were treated with PXS-5505 for 7 days and gene expression quantified by qRT-PCR. **(H)** Schematic showing how the deposition and sensing of matrix components controls key transcriptional mediators in ILC, and the perturbance of this signalling axis upon LOX inhibition. Statistical significance determined by Student t-test or one-way ANOVA. * P < 0.05, ** P < 0.01, *** P < 0.001.

Interrogation of the TNFα Signalling via NFκβ gene set revealed coordinated downregulation of key transcription factors *MYC, JUN* and *FOS*, as well *CCN1,* encoding a matricellular protein that promotes cell-ECM adhesion and binds to integrins including the *ITGAV* and *ITGB5* (Fig. 7B). CCN1, regulated by the AP-1 complex (61, 62), initiates signalling cascades through integrin-FAK engagement, culminating in NF-κB activation and transcription of genes that reinforce cell survival, adhesion, and MYC/ AP-1 activity (63, 64).

Expression of these 4 genes was significantly increased in ILC relative to ER+ NST tumours (TCGA) (Fig. 7C), demonstrating the relevance of our observations on CCN1/MYC/Jun and FOS signalling to clinical samples. The TNFα Signalling via NFκβ signature was enriched to a lesser extent in ILCs vs NSTs (Supp Fig. G). Furthermore, MYC Targets _V1 or V2 hallmark pathways were consistently down regulated by LOXi in the SUM44, MM134 and IPH-926 models (Supp Fig. D). In the SUM44 model, cell cycle activity at protein level (REACTOME, Fig. 5D) was decreased across all treatments, yet, transcriptional suppression of MYC_TARGETS_V1 was specifically observed in the OVX+PXS combination group, suggesting that LOXi may confer additional, repression of transcriptional MYC activity not fully captured by protein-level assays.

To further understand the additional effects of LOXi we performed a differential expression analysis on proteins down modulated by the combined therapy but not standard of care (OVX). This identified 108 proteins in the SUM44 and 220 in the MM134 model (Fig. 7D). Enrichment analysis of this protein subset revealed significant suppression of MYC target pathways. Analysis of ChIP-seq data (60) confirmed MYC as the top enriched transcription factor target regulating these proteins (SUM44, adj. p-value=2.16^-8^, MM134 adj. p-value=1.582^-15^) (Fig. 7E). In the METS27 model, enrichment analysis on downregulated proteins also identified

MYC targets as the most strongly downregulated pathway (Supp. Fig. 5D,E). These findings suggest that MYC signalling may serve as a surrogate biomarker of LOXi activity detectable at both the proteomic and transcriptomic levels. Finally, we asked whether matrix stiffness alone could regulate these transcriptional dependencies. ILC cells were cultured in type 1 collagen matrices of 1 or 4mg/ml concentration, corresponding to physiologic and tumour-associated stiffness levels, respectively (54). Higher matrix density induced expression of *LOXL1* as well as *MYC, JUN*, *FOS* and *CCN1* transcripts (Fig. 7F). When cells cultured under stiff (4mg/ml) conditions were treated with PXS-5505 for 7 days, transcript expression of *JUN*, *FOS* was significantly decreased, whilst *CCN1* and *MYC,* showed a trend (Fig. 7G). These findings support a role of collagen density and LOX activity in regulating these transcription factors.

Taken together, these findings illuminate a critical ECM-cell adhesion-transcriptional axis in ILC, wherein collagen crosslinking and matrix stiffness sustain integrin-mediated MYC/AP-1 transcriptional programs which include CCN1 as downstream target and upstream regulator. LOX inhibition disrupts this axis - impairing ECM architecture, integrin signalling, and MYC/AP-1-driven gene expression resulting in further decrease in ITG expression and ultimately decreased cell proliferation (Fig. 7H). New clinical readouts, including imaging and immunohistochemistry, as well as proteomic and transcriptomic analyses converge to support the rationale for a LOX-targeting clinical trial in ILC.

## Discussion

Through integration of unbiased synthetic lethality screens and intraductal xenograft models we have unveiled a unique vulnerability of CDH1 deficient ILCs related to compensatory integrin signalling, characterised by increased expression and activity of the transcription factors MYC and AP-1, which operate within in a positive feedback loop involving ITGs and CCN1. While the powerful genetic screens conducted in 2D cultures identified critical cell-intrinsic components required for fitness in the absence of *CDH1*, the *in vivo* intraductal models allowed us to unravel essential extracellular components upstream of the integrin-MYC axis that were not detected *in vitro*. These components include at least two elements: biophysical matrix properties and ECM/cell surface complexes.

The MIND models employed in our study reflect both intra-patient and inter-tumoral heterogeneity at the transcriptomic and histopathological levels during disease progression and treatment responses, bringing preclinical research closer to translation. Our multiparametric analysis of drug response recapitulates the difficulties in quantifying ILC disease extent clinically. We show that collagen fiber architecture and gene expression changes are more sensitive, robust, and may serve as surrogate markers to enable clinical trials. To identify predictive biomarkers, such as *CDH1* levels, it will be useful to compare transcriptomic profiles of responders and non-responders. For this, larger sets of models or clinical trials are needed.

Our finding that *MYC, FOS* and *JUN* are upregulated in ILCs compared to ER+ non ILC tumours in patient datasets strongly reinforces the clinical relevance of both our results and models. Consequently, incorporating morphological features as an additional dimension to consider when deciding about treatments, may add granularity and improve therapeutic precision. Artificial intelligence (AI)-based approaches are expected to lead to advances in this direction.

The transcription factor JUN in particular, is highly expressed in fibrotic tissues characterised by abnormal collagen deposition (65). It’s overexpression also reprograms ER chromatin binding and reduces tamoxifen sensitivity (66), while stiff ECM can enhance the agonistic and pro-proliferative effects of tamoxifen (67). This raises the intriguing possibility that the clinically observed reduced sensitivity to tamoxifen of ILCs (68) may be influenced by matrix stiffening and JUN mediated effects. Collectively, these observations provide a strong rationale for assessing a LOXis in combination with endocrine therapy in ER+ ILC.

While our window of opportunity trial model focuses on early-stage disease, PXS-5505 may also benefit patients with late-stage disease. Of note, LOX inhibition did not produce a detectable reduction in tumour growth in the ER- metastatic IPH-926 model at a late stage of disease but did impact metastatic dissemination, in line with the role of LOX in priming the pre-metastatic niche (69). This may extend to other models if allowed to progress to the metastatic stage. Additionally, extending LOXi treatment to two months in the aggressive ER+ METS27 model slowed primary tumour growth, suggesting potential benefits of longer treatment durations in advanced-stage disease.

LOXis have demonstrated antitumor effects by targeting cancer-associated fibroblasts (CAFs) and reducing fibrosis in desmoplastic pancreatic cancer (28), and TNBC xenograft models (16). The present study unveils a different mechanism whereby, in the case of ILCs, LOXi exerts direct effects on tumour cells through the loss of integrin signalling by disrupting collagen fibers, resulting in decreased cell cycling and cell loss.

Combining LOXi with chemo- or immunotherapy could further enhance treatment efficacy. LOX activity in T- cells within the lungs increased in response to paclitaxel therapy, accompanied by heightened ECM remodelling and metastasis in preclinical breast cancer models (70), with this effect being mitigated by LOX inhibition using BAPN. Notably, LOXi have been shown to modulate the ECM in ways that improve T-cell migration and enhance the efficacy of PD-L1 blockade—a particularly promising avenue for addressing immune "cold" phenotypes observed in a subset of ILCs (71).

One limitation of our study is the immunocompromised setting necessitated by PDX models. Although it is not possible to assess immune responses in this setting, efforts are ongoing to develop murine models of ILC that will allow these questions to be addressed (7). Intraductal xenografts also show significantly less CAF infiltration, but do show macrophage infiltration (72), and it cannot be ruled out that the effects of LOXi may also be mediated through these cell types.

Taken together, we show that PXS-5505, a highly selective inhibitor with an excellent pharmacodynamic profile well tolerated in pre-clinical and clinical settings, is an excellent candidate either as a monotherapy or in combination with standard-of-care AI in ILC patients. The models used and endpoints identified in this study provide an entry point for clinical trials in this underexplored space.

## Methods

### Data pre-processing

SCAN-B and TCGA bulk RNA-sequencing (RNA-seq) datasets were downloaded and pre-processed as previously described (22). The neoadjuvant chemotherapies (NAC) RNA-seq transcript per million (TPM) dataset was downloaded from GEO using the accession code GSE123845.

### Data normalisation

All 3 datasets were normalised using EMBER’s procedure (22). In short, for each sample individually, genes are ranked from lowest to highest expression, rankings are divided by an average ranking of stable genes and each sample is embedded in the EMBER space. The embedding removes batch effects and allows direct comparison of signature scores across datasets.

### Survival analysis

Standardised survival probabilities were calculated using the R package flexsurv (v2.3) (73). The median of either the genes or the gene signatures was used as a threshold to dichotomise the variables and standardised survival probabilities were adjusted by age and tumour stage.

### Cell culture

Cell lines were obtained from ATCC and maintained at 37°C in humidified incubator in an atmosphere of 5% CO2. MDA-MB-134IV (MM134), SUM44PE and IPH-926 expressing RFP-Luc2 were cultured as previously described (8, 12, 36). Cells were plated at a density of 2x10^4/well of a 96 well, or at 5x10^4/well in 1 or 4mg/ml collagen matrices (Rat Tail Collagen Type 1, Gibco). Cells were treated with DMSO, PXS-5505 (10 µM) BAPN (10µM) or Fulvestrant (100nM) Media containing compounds was refreshed on day 2 and day 5. For RNAi experiments cells were plated in 96 well plates in triplicate per condition, and transfected with RNAiMAX (3ul/ well) and siRNA or a non-targeting scrambled control (20nM) (Qiagen) in Opti-MEM. Plates were imaged (Brightfield and red fluorescence) at 6hr intervals using an Incucyte (Sartorius).

### Synthetic lethal analysis

Genome-wide CRISPR-Cas9 screen data, transcriptomic data and mesenchymal/epithelial classifications for tumour cell lines was accessed from depmap.org/portal/ (47) on 21 March 2025. E-cadherin high/low expression cohorts of tumour cell lines were defined using CDH1 mRNA expression (as shown in Figure 6B) and then used to identify synthetic lethal genes (Figure 6C) using the depmap.org/portal/ (47) custom analysis tool. Mesenchymal vs. epithelial dependencies were also identified using the depmap.org/portal/ (47) custom analysis tool.

### Clinical samples/PDX derivation

In the UK written informed consent was obtained in accordance with the Declaration of Helsinki under the research ethics committee approved studies (REC reference14/LO/0292, Royal Marsden NHS Foundation Trust, UK). In Switzerland the study involving human subjects was approved by the Commission cantonale d’éthique de la recherche sur l’être humain **(CER-VD 38/15; PB_2016-01185)**, and informed consent was obtained for all human subjects participating in this study.

For pleural effusions the sample was centrifugated 1200 rpm for 15 minutes. Cell pellet was resuspended in red blood cell lysis buffer Hybri-Max (Sigma) and incubated at room temperature for 20 minutes. Cells were washed in L15 medium and resuspended in L15 medium (10% FCS and Primocin) and seeded in a tissue culture flask for 3-4 hours after which the floating tumour cells were collected frozen down for *in vivo* and *in vitro* assays.

Solid tumour tissue was mechanically and enzymatically digested using parallel razor blades and collagenase, as previously described (30, 74). Pellets were rinsed with phosphate-buffered saline (PBS), 2% calf serum (CS), and erythrocyte-lysed using red blood cell lysis buffer (Sigma, R 5) for 5 min, then diluted in PBS 2% CS, and centrifuged again. Patient-derived tumour cells were transduced with either ffLuc2/Turbo-GFP or ffLuc2/Turbo-RFP (GEG-tech) overnight in low attachment culture plates (Corning® Costar® Ultra-Low Attachment) in a humidified incubator (3 °C, 5% CO2, and 5% O2).

### *In vivo* experiments

All animal studies were conducted in accordance with protocols approved by the Institute of Cancer Research, London, UK and with the UK Animals (Scientific Procedures) Act 1986. Studies carried out at EPFL were approved by Service de la Consommation et des Affaires Vétérinaires of the Canton of Vaud, Switzerland. NSG (NOD.Cg-PrkdcSCID Il2rgtm Wjl/S J) mice (Charles River, Harlow, K) were maintained in a 2-h-light- 2-h- dark cycle, with controlled temperature and food and water ad libitum. Eight- to 2-week-old female mice were anesthetised by isoflurane and intraductally injected into the th mammary glands with μl of PBS containing 200,000–5, cells. Luciferase-based imaging was performed with Xenogen I IS Imaging System 200 (Caliper Life Sciences) in accordance with the manufacturer’s protocols and used to monitor individual mammary glands. At sacrifice, engrafted mammary glands and organs were harvested, fixed in 4% paraformaldehyde for histology and IHC or snap-fro en for RNA and protein isolation. Stereo-micrographs of engrafted glands were acquired with a THUNDER Imager Model Organism (Leica).

For the pharmacologic inhibition of LOXLs, mice with established tumours as determine by BLI were randomised to receive a diet containing vehicle or PXS-5505 (Research Diets) for 28 days, with or without ovariectomy. Surgery was performed 7 days prior to the addition of compound-containing diet and was carried out under anaesthesia as previously described, with non-ovariectomised mice receiving a sham surgery. For shorter treatment windows PXS-5505 (LOX-IN-3, Medchem Express) was injected intraperitoneally at a daily dose of 30 mg/kg for 14-21 days. BLI growth curve analysis was performed using BioGrowler, a Bayesian hierarchical modelling tool (35).

### Digital pathology

Computational pathology analysis was performed using a previously published dataset (24).

### Immunohistochemistry

All samples were fixed in % paraformaldehyde and paraffin-embedded (FFPE). For immunostaining, 4-μm sections were mounted onto 76 × 26 mm microscope slides. Hematoxylin and eosin was performed according to standard protocols. PSR was performed using a lab-validated assay using the a ready-to-use PSR solution (AbCam, ab236842) and proprietary two-part Weigert’s Iron Haematoxylin kit (Merck, 1.15973)

Immunostaining for Ki67 was performed using anti-Ki67 MIB1 was on the Dako/Agilent Link48 Autostainer CE- IVD automated immunostaining platform. Prior to staining, heat-induced epitope retrieval and simultaneous dewax was performed using the Dako/Agilent PT-Link module using pH6 Target Retrieval solution (Agilent Technologies, K800521-2) for 20mins at 97c according to manufacturers instructions. All subsequent incubations were carried out at room temperature and rinses performed with Dako Wash Buffer (Agilent, K800721-2). Endogenous peroxidases were blocked for 5 minutes using Dako REAL Peroxidase Block (Agilent, S202386-2) then primary antibody (Ki67 mouse anti-human monoclonal MIB1, Agilent Technologies, M724001-2) diluted 1/300 in FLEX Primary Antibody Diluent (Agilent, K800621-2) applied for 60 minutes. Bound antibody was detected using Dako Anti-Mouse EnVision Polymer-HRP (Agilent, K400111-2) and the reaction visualised using Dako DAB+ (Agilent, K346811-2). Nuclei were counterstained using Dako FLEX Haematoxylin (Agilent, K800821-2). Whole slide digital images were captured on a Hamamatsu Nanozoomer at 20x magnification.

Tumour deconvolution was performed using a trained classifier on QuPath to segment tumour, stroma and adipose. Ki67 scoring was performed on QuPath taking <5 regions of interested (ROI’s) around ducts containing tumour cells and expressed as % positive cells.

### Collagen morphometrics analysis

At least five regions of interested (ROI’s) containing tumour cells were selected from each xenografted mammary gland. Colour deconvolution was performed using ImageJ to separate the fibrillar collagen stain from the cellular counterstain. Images were processed using the TWOMBLI analysis pipeline as published previously and expressed as average value per tumour. Correlation matrices were generated using collagen morphometric outputs, % Ki67 expression and fold change total flux at end point for each tumour.

### RNA extraction from cells

Cells were extracted from collagen matrices by the addition of 100 uL of 0.1% collagenase (Roche, UK) to each well. Cells were collected in a pellet and after aspirating the supernatant, the RNA was extracted by following the standardised protocol for non-adherent cells using the Bio-Rad SingleShot Cell Lysis Kit. For 2D cultures the adherent cell protocol was followed. cDNA was synthesised using the ThermoFisher High Capacity cDNA Reverse Transcription Kit as per the manufacturer’s instructions.

### RNA extraction from tissues

Mammary glands bearing tumours were homogenised at 4°C in 700ul QIAzol lysis buffer in 2ml screw cap tubes with metal beads in a Precellys tissue homogeniser. Lysates were precipitated with chloroform and purified using RNeasy kits (Qiagen, UK) on a Qiacube. RNA concentrations and purity were determined using a Nanodrop Spectrophotometer.

### qPCR

A Power p SYBR Green (Applied Biosystems) master mix was prepared according to the manufacturer’s instructions using Quantitect Primer Assay for human *GAPDH, COL1A1, LOXL1, ITGB5, ITGAV, JUN, MYC* and *MKI67* (Qiagen, UK) and 5-10ng of cDNA. Samples were run in duplicate and Ct values were obtained using a LightCycler (Roche). Change in expression was measured using the ΔΔCt method and expressed as relative expression versus the experimental control or an internal universal reference (Thermo Fisher).

### Proteomics

Tissues were homogenised in 2ml screw cap tubes with metal beads in the Precellys system using 1mL of lysis buffer containing 1% sodium deoxycholate (SDC), 100 mM triethylammonium bicarbonate (TEAB), 10% isopropanol, 50 mM NaCl freshly supplemented with Halt protease and phosphatase inhibitor cocktail (100X) (Thermo, #78442). The homogenised samples were transferred into 2 mL Eppendorf tubes with a wide edge tip and probe sonicated for 1 min with pulses of 1 sec on ice (EpiShear, amplitude 40%). Samples were clarified with centrifugation at 13,000 rpm for 10 min and protein concentration was measured with the Quick Start Bradford protein assay (Bio-Rad). Aliquots of 5 μg of total protein were reduced with 5 mM tris-2- carboxyethyl phosphine (TCEP), alkylated with 10 mM iodoacetamide (IAA) and digested overnight with trypsin (Pierce, ratio 1:20) at room temperature. Peptides were labelled with the TMTpro multiplex reagents (Thermo) according to manufacturer’s instructions. The peptide pool was fractionated with high pH Reversed- Phase (RP) chromatography using the XBridge C column (2. x 5 mm, 3.5 μm, Waters) on an ltiMate 3000 HPLC system over a 1% gradient in 35 min. Mobile phase A was 0.1% (v/v) ammonium hydroxide and mobile phase B was 0.1% ammonium hydroxide (v/v) in acetonitrile. LC-MS analysis was performed on an UltiMate 3000 system coupled with the Orbitrap Ascend Mass Spectrometer (Thermo) using a 25 cm capillary column (Waters, nanoE MZ PST BEH 3 C, . μm, 5 μm × 25 mm) over a min gradient 5%-35% mobile phase B composed of 80% acetonitrile, 0.1% formic acid. MS spectra were collected at Orbitrap mass resolution of 120k and precursors were targeted for HCD fragmentation in the top speed mode (3 sec) with collision energy 32% and iontrap detection in turbo scan rate. MS3 scans were triggered by Real Time Search (RTS) against a Fasta file containing UniProt Homo sapiens and Mus musculus reviewed canonical sequences with multi-notch isolation (10 notches) and HCD fragmentation with collision energy 55% at 45k Orbitrap resolution. Targeted precursors were dynamically excluded for further activation for 45 seconds and RTS close- out was enabled with max 4 peptides per protein. The Sequest HT and Comet nodes in Proteome Discoverer 3.0 (Thermo) were used to search the raw mass spectra against the same Fasta file that was used for RTS acquisition. The precursor mass tolerance was set at 20 ppm and the fragment ion mass tolerance at 0.5 Da (or 1 Da for Comet). TMTpro at N-terminus/K and Carbamidomethyl at C were defined as static modifications. Dynamic modifications were oxidation of M and deamidation of N/Q. Peptide confidence was estimated with the Percolator node and peptide FDR was set at 0.01 based on target-decoy strategy. Only unique peptides were used for quantification, considering protein groups for peptide uniqueness. Peptides with average reporter signal-to-noise greater than 3 were used for protein quantification.

Normalised log2 median-centred values were used for differential analysis. Proteins that reach a threshold of p<0.05, logFC >0.5,-0.5 were considered significantly differentially expressed. Pathway enrichment was performed on significantly up or downregulated protein lists using EnrichR (60) and STRING (75). Matrisome AnalyzeR (42) was using to annotate matrisomal and associated human and mouse proteins.

### Transcriptome profiling

RNA-Seq profiling generated 35.9 to 60.1 million paired-end reads per sample. Prior to alignment, MIND RNA- Seq FASTQ reads were analysed by the BBMap (v38.87) BBSplit function to remove mouse aligning reads (cite BBSplit - Bushnell B. - sourceforge.net/projects/bbsplit/). FASTQ reads were next trimmed using Trim Galore (v0.6.6). FastQC, FastQ Screen (76) and MultiQC (v1.9) (77) were ran to determine library quality. Paired-end reads (150bp) were aligned to the human reference genome GRCh38 using STAR 2.7.6a (78) with --quantMode GeneCounts and --twopassMode Basic alignment settings, against annotation files downloaded from GENCODE (v22) in GTF file format. Results were further annotated using ENSEMBL gene annotations with the R package org.Hs.eg.db (v3.10.0) in the R statistical programming environment (v3.6.0). Differential gene expression analysis was performed using edgeR (v3.28.1) (79) with limma (v3.42.2). Genes with low expression were filtered out using edgeR’s function ‘filterByExpr()’ with default parameters. Raw counts were normalised using edgeR’s TMM (trimmed mean of M-values) method and differential mRNA abundance was performed using the quasi-likelihood (QL) F-test. sing edgeR’s glmQLFTest statistics, genes were ranked as: –log_10_(P) x sgn(log_2_ fold-change) and analysed with the Fast Gene Set Enrichment Analysis (FGSEA) R package fgsea (v. .), R (v . .). FGSEA was performed against all “hallmark” gene sets (v .5.).

### Statistical analysis

Statistical analysis was performed using Graphpad Prism unless stated otherwise. Comparisons between groups of continuous variables were made using an unpaired two-tailed Student’s *t*-test or ANOVA. All tests were two-sided and a *P* value of less than 0.05 was considered significant. Data points represent biological replicates, and for in vivo experiment the number of individual tumours (glands) is detailed in the figure or legend.

## Data availability

Mass spectrometry proteomics data have been deposited to the ProteomeXchange Consortium via the PRIDE60 partner repository with the dataset identifier PXD065164.

ranscriptomic data is deposited under ascension PRJNA1284888. <u>List of Supplementary Materials</u>

Materials and Methods Supplementary Figures S1 to S6

## Acknowledgements and Funding

We thank Breast Cancer Now for funding this work as part of Programme Funding to the Breast Cancer Now Toby Robins Research Centre. Sanz- Moreno lab was further supported by The Institute of Cancer Research, Cancer Research UK (CRUK) C33043/A24478; Barts Charity; World Wide Cancer Research 22-0329 and UKRI grant reference EP/X033392. Y.L. acknowledges the support of the China Scholarship Council program (Project ID: 202306100230). We would also like to thank Syntara for their support in providing experimental compounds, as well as the Histopathology Core Facility and thank Breast Cancer Now for supporting the work of this team. Finally, we thank the patients who contributed material to the generation of models in this study.

## Author Contributions

Conceptualization: RLF, GS, CB Methodology: RF, FH, GS, CR, HK, TH, GA, LB, HP, SP, SJ, LP, WJ, AI, AS, KZ, MF, JSC Investigation: RF, FH, GS, AA, AZ, SB, YZ, HMQ, YL, RM, MI, BAH Visualization: CR, HK, TR Funding acquisition: CB, VSM, NCT, CJL, CMI Writing – original draft: RF, CB Writing – review & editing: RF, CB

