## Supplementary material for "LOX inhibition disrupts a collagen-integrin–MYC axis as a translatable targeting strategy in invasive lobular carcinoma": Supplemenary figures

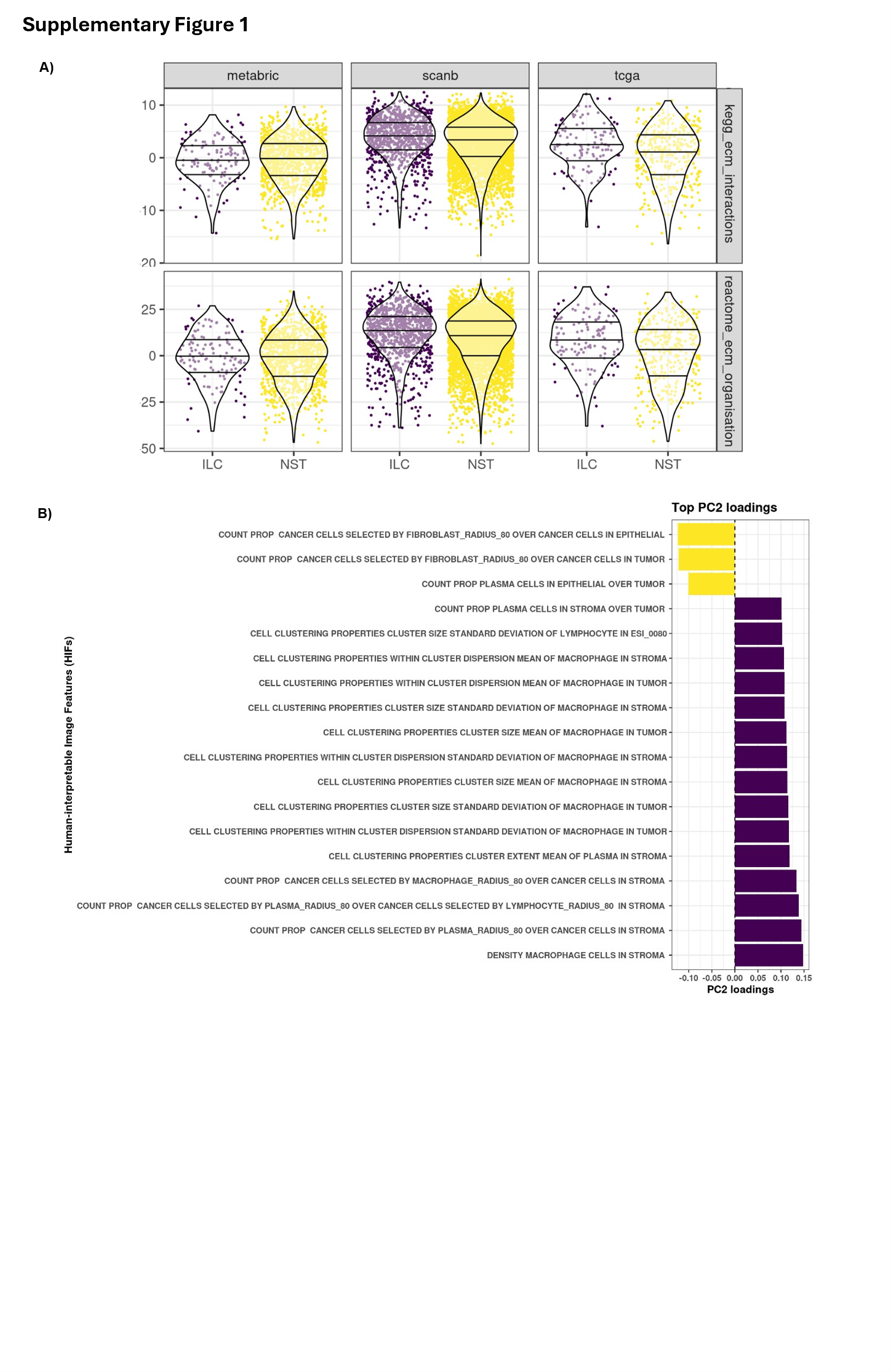

***Supp Fig. 1 – (A)*** *Violin plots showing regressed matrisome signature score in ILC vs non-ILC (ER+ NST) in TCGA, METABRIC and SCAN-B cohorts****. (B)*** *Principle component loadings for PCA plot shown in Fig. 1E.*

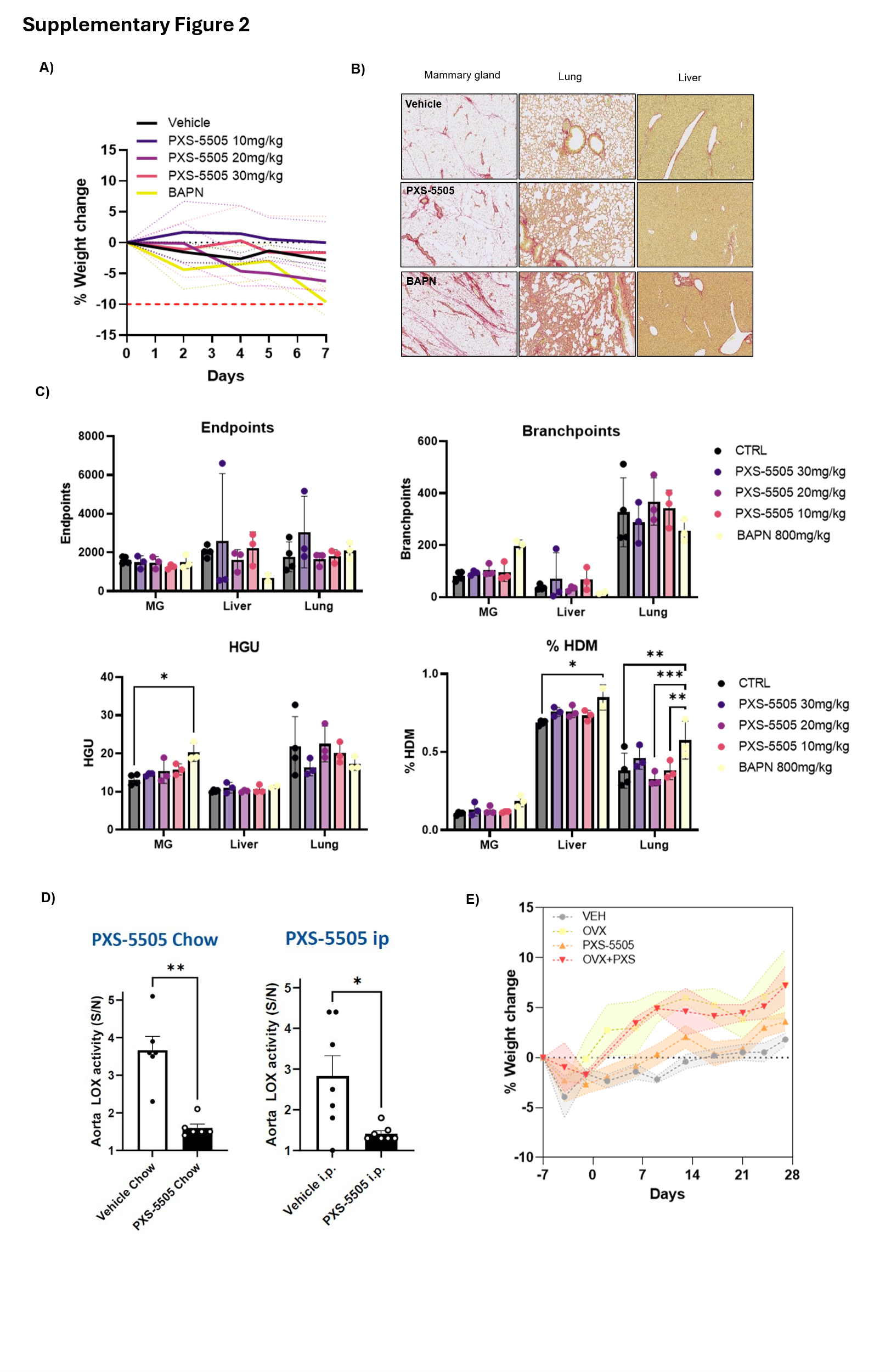
***Supp Fig. 2 (A)*** *Weight change of healthy female NSG mice treated with PXS-5505 (daily i.p, 10-30mg/kg) or BAPN (800mg/kg) for 7 days.* ***(B-C)*** *Quantification of collagen morphometrics in the mammary gland, lungs and liver.* ***(D)*** *LOX activity in the aorta of mice exposed to PXS-5505 by i.p or in the diet.* ***(E)*** *Weight change of* *intraductal xenograft bearing mice having underwent ovariectomy or sham surgery and given vehicle or PXS-5505 diet.*

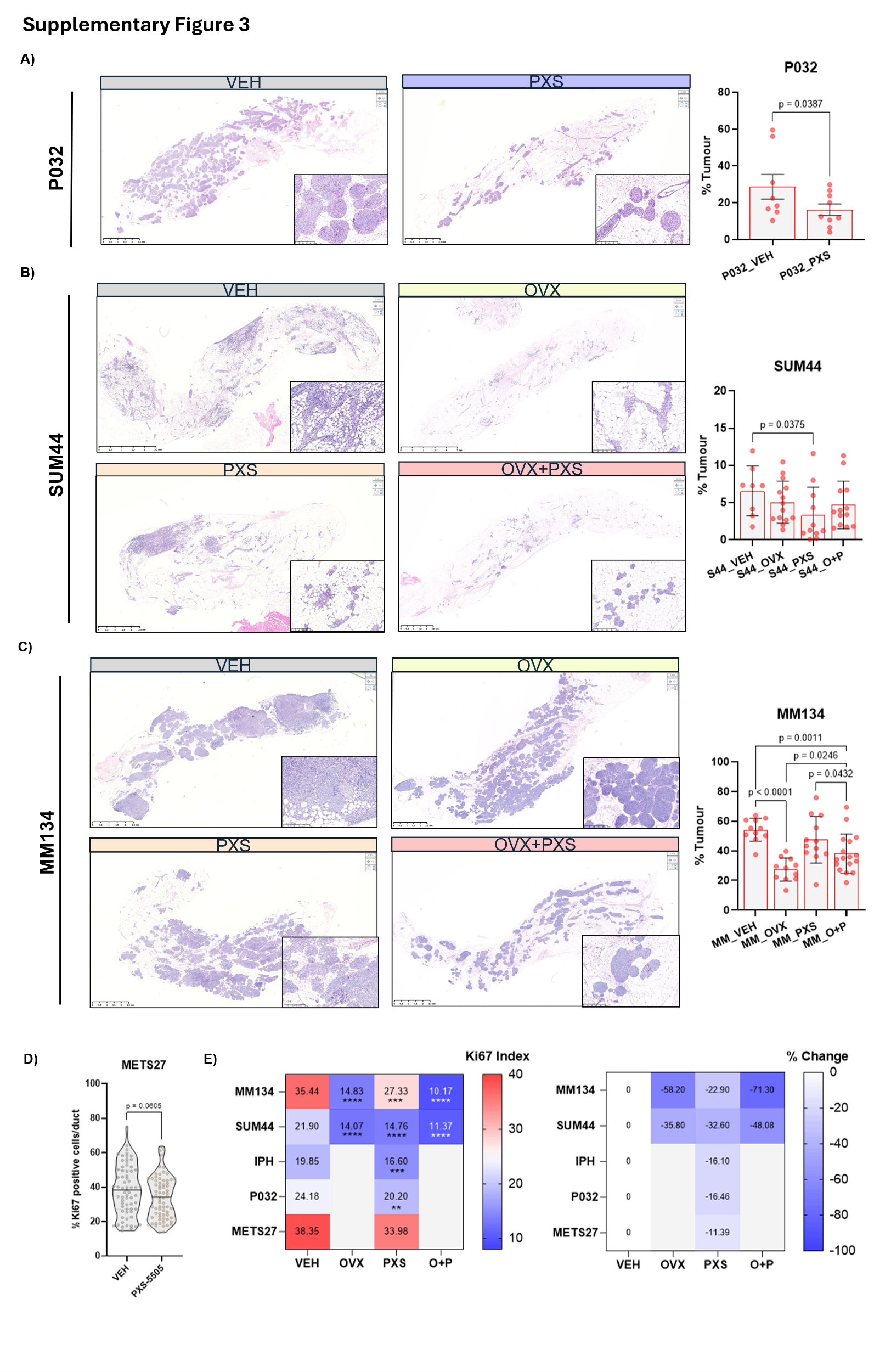

***Supp Fig. 3 – (A-C)*** *Representative H+E stained sections from intraductally engrafted glands from P032 (top) SUM44 (middle) and MM134 (bottom) models (left). Histopathological composition analysis of intraductal xenografts at endpoint (right).* ***(D)*** *Violin plot showing % Ki67 positive tumour cells/duct quantified by IHC in METS27.* ***(E)*** *Heatmap of average Ki67 indices across the intraductal models tested (left), and expressed as % change compared to vehicle (right).*

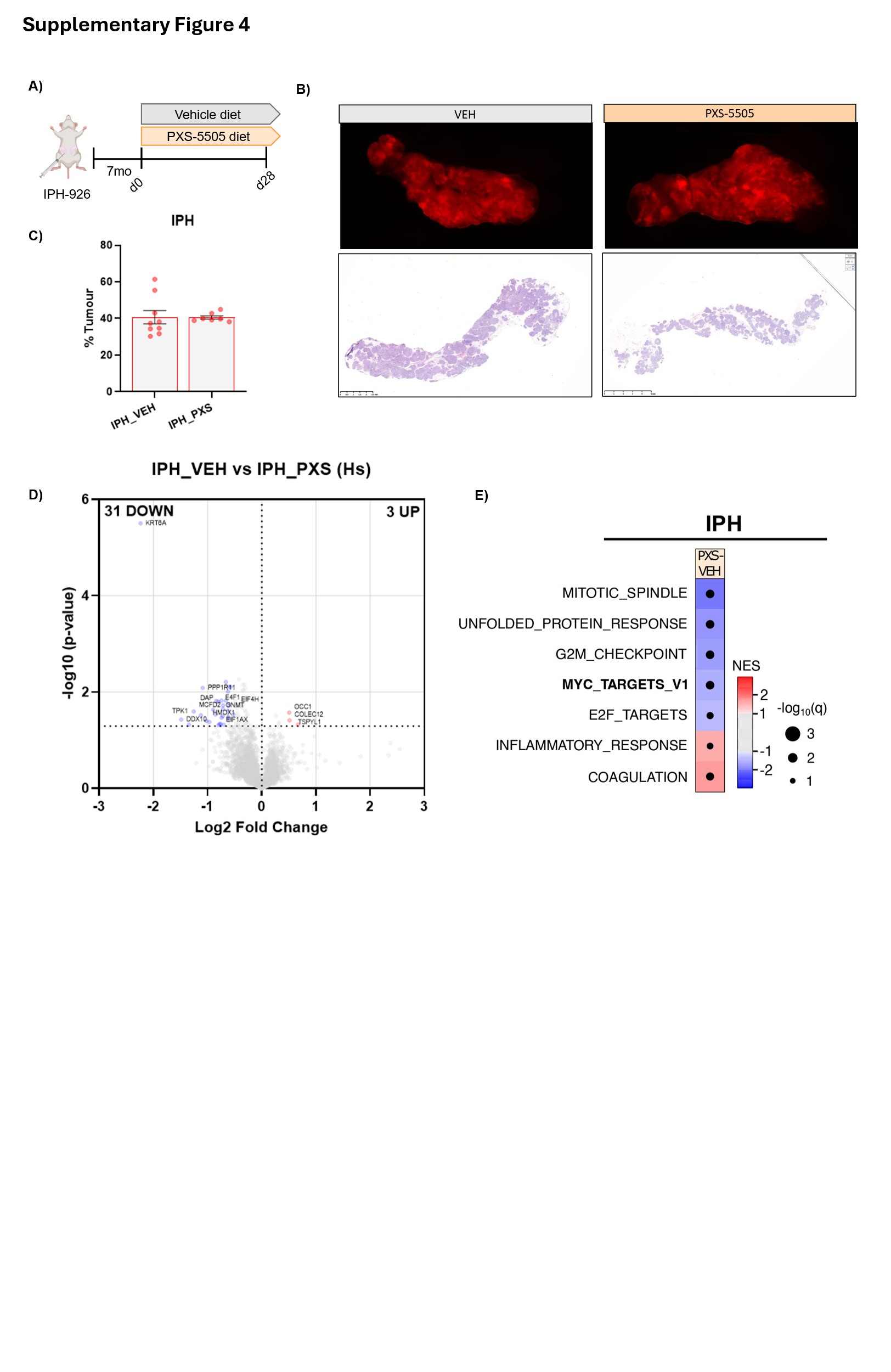
***Supp Fig. 4 – (A)*** *Experimental scheme* ***(B)*** *Representative fluorescence stereomicrographs of IPH-926 engrafted inguinal glands shown (top), and H+E stained sections (bottom).* ***(C)*** *Histopathological composition analysis of intraductal xenografts at endpoint (right).* ***(D)****Volcano plot of differentially regulated proteins in Vehicle vs PXS-5505 treated primary tumours.* ***(E)*** *Heatmap showing* *normalised expression score (NES) values for HALLMARK gene sets from RNAseq profiling of PXS-5505 treated mice vs Vehicle.*

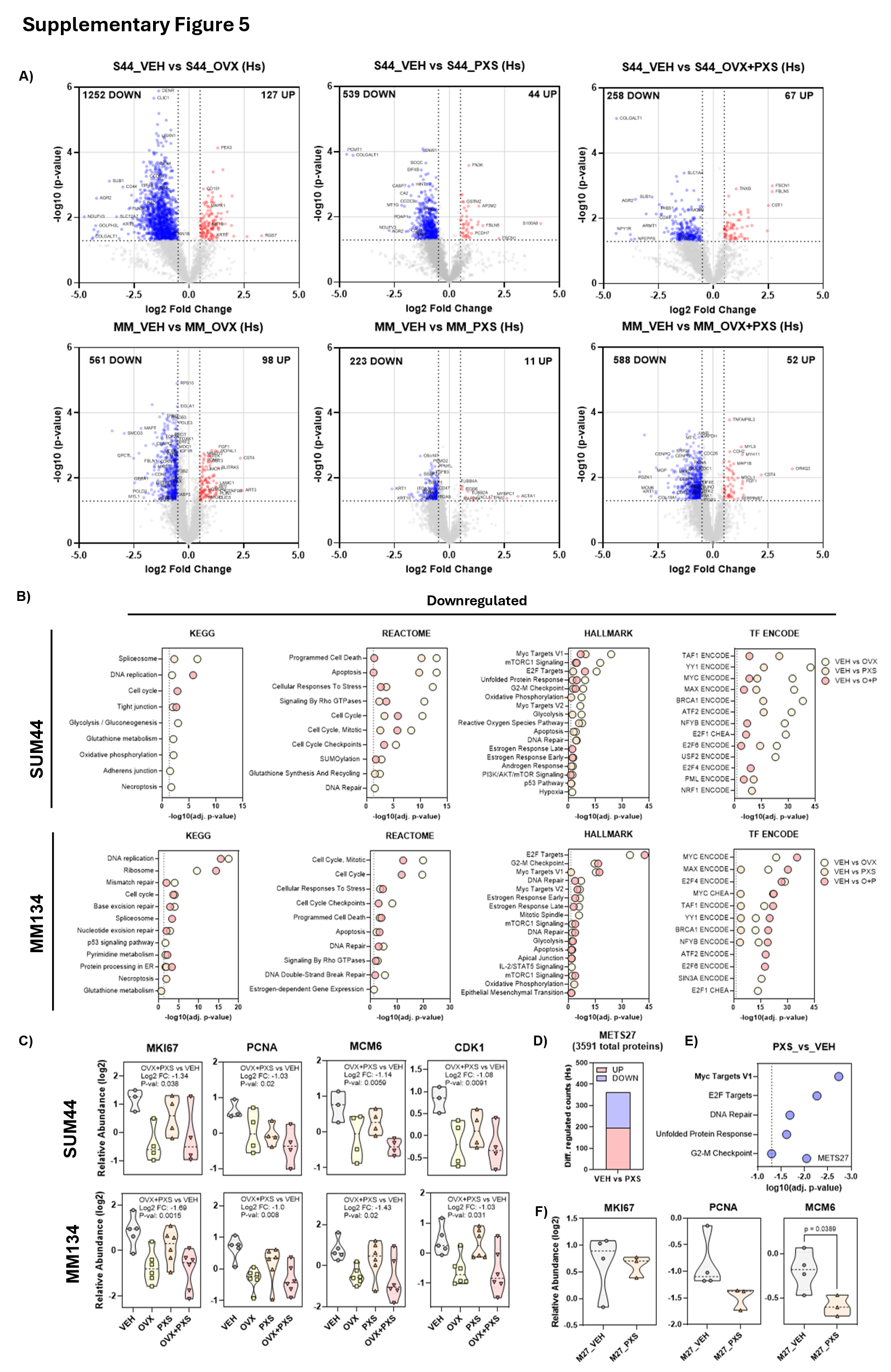

***Supp Fig. 5– (A)*** *Volcano plots of differentially regulated proteins across each condition in SUM44 (top) and MM134 (bottom) models.* ***(B)*** *Pathway enrichment for downmodulated proteins across treatment conditions vs vehicle.* ***(C)*** *Protein abundance of proliferation and cell cycle markers.* ***(D)*** *Number of proteins differentially upon PXS-5505 treatment in the METS27 model.* ***(E)*** *Pathway enrichment of downregulated proteins.* ***(F)*** *Protein abundance of proliferation and cell cycle markers.*

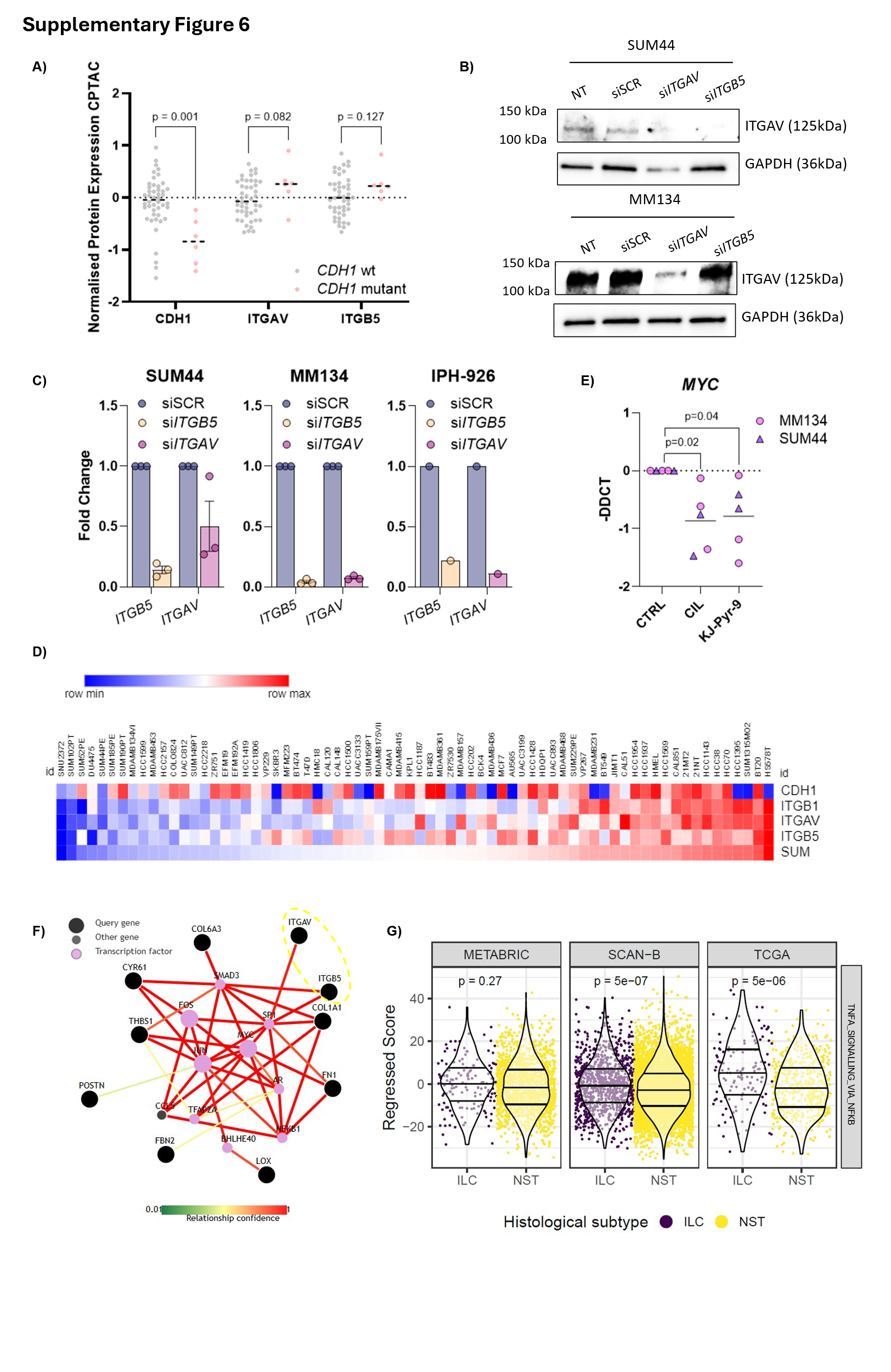
***Supp Fig. 6 – (A)*** *Mass spectrometry proteomic data from 52 ER+ breast cancers collated in the TCGA study. Data was accessed 21 June 2025. Normalised Protein Protein expression levels of CDH1 (E-cadherin), ITGB5 and ITGAV are shown in E-cadherin mutant (CDH1 mutant) and E-cadherin wild type tumours. In this case CDH1 mutant cancers were defined as those with deleterious truncating mutations. P values calculated by Mann Whitney t test.* ***(B)*** *Protein expression of ITGAV in siRNA transfected cells.* ***(C)*** *Bar graphs of ITGAV and ITGB5 expression determined by qRT-PCR in ILC cell lines transfected with siRNA directed against ITGAV and ITGB5 as well as a scrambled control (siSCR).* ***(D)*** *Heatmap of log2 TPM expression of CDH1, ITGB1, ITGAV, ITGB5 and the sum integrin expression in breast cancer cell lines from the Cancer Cell Line Encyclopaedia database.* ***(E)*** *Expression of MYC determined by qRT-PCR in ILC cells in 4mg/ml collagen matrices exposed to CIL and 20µM KJ-Pyr-9.* ***(F)*** *PathwayNet map of genes and transcription factors relating to proteins identified as downregulated by LOXi.* ***(G)*** *Violin plot showing regressed TNFA_SIGNALLING_VIA_NFKB score in ILC vs non-ILC (ER+ NST) in TCGA, METABRIC and SCAN-B cohorts.*
